# A *Saccharomyces boulardii* synthetic biotic platform for delivery of therapeutic nanobodies to ameliorate gastrointestinal inflammation

**DOI:** 10.1101/2025.05.15.652971

**Authors:** Roger Palou, Almer M. van der Sloot, Aline A. Fiebig, Megan T. Zangara, Naseer Sangwan, María Sánchez Osuna, Bushra Ilyas, Haley Zubyk, Michael Cook, Gerard D. Wright, Brian K. Coombes, Mike Tyers

## Abstract

Protein-based pharmaceuticals, such as engineered antibodies, form a major drug class of steadily increasing market share. However, these biologic medicines are costly to manufacture, are subject to strict supply chain and storage constraints and often require invasive administration routes. Engineered microbes that secrete bioactive products directly within the microbiome milieu may mitigate these challenges. Here, we describe a cell microfactory platform based on the probiotic yeast *Saccharomyces boulardii* for the production of nanobody biologics in the gastrointestinal (GI) tract. High-level secretion of nanobodies by *S. boulardii* was achieved by optimizing promoters, secretion signals, and antibody formats. In mice, oral gavage of *S. boulardii* allowed efficient and transient colonization of the colonic compartment and in situ production of a therapeutic nanobody directed against tumor necrosis factor (TNF). In a mouse model of chemical-induced colitis, GI-delivery of anti-mTNF nanobody via live *S. boulardii* improved both survival and disease severity without causing overt perturbation of microbiome composition. These results position *S. boulardii* as a synthetic biotic platform for the in situ production and delivery of protein-based therapeutics to the GI tract.

## Introduction

The concept of using live microbial cells to manufacture and deliver therapeutic agents to the microbiome – variously termed synthetic biotics, engineered probiotics, cell microfactories or living therapeutics – holds promise for cost-effective, site-specific therapeutic intervention^1,2^. Synthetic biotics may be designed to produce biosynthesized small molecules such as nutrients, vitamins and antibiotics, or more complex protein-based agents such as antibodies, peptides, growth factors and cytokines. The in-situ synthesis of active biomolecules in the gastrointestinal (GI) tract by cell microfactories is now well-established for bacterial platforms, including *Lactobacillus lactis*, *Lactococcus lactis* and *E. coli* Nissle 1917^3–6^. However, the use of bacterial species as orally administered synthetic biotics can be compromised by loss of viability in the acidic environment of the stomach, the inability to produce high levels of secreted complex biologic molecules such as antibodies, and the risk of lateral gene transfer to other species in the microbiome or external environment^3,7^. Although the gut microbiome is dominated by bacterial species, fungal, archaeal and protozoal species form another important component of the microbiome^8^.

To overcome the problems inherent to bacterial-based synthetic biotics, the yeast *Saccharomyces boulardii* has been explored as an alternative platform for in situ production of biologic agents ^9–13^. *S. boulardii* is closely related to the budding yeast *Saccharomyces cerevisiae*^14^, also known as baker’s or brewer’s yeast, a mainstay model organism for eukaryotic molecular genetics that is also widely used in food and biotechnology production processes. *S. boulardii* is a generally recognized as safe (GRAS) organism that is widely used as an over-the-counter probiotic for the treatment of *Clostridium difficile* infections and antibiotic-associated diarrhea^15^. *S. boulardii* has inherent potential advantages as a synthetic biotic that include relative ease of genetic manipulation that benefits from extensive molecular genetic tools developed over many decades for *S. cerevisiae*. Like budding yeast, *S. boulardii* can be readily produced at an industrial scale and at low cost, retains long term viability in desiccated form, and shows negligible rates of lateral gene transfer^15,16^. Moreover, *S. cerevisiae* spp. are part of the normal human mycobiome^17^. Unlike *S. cerevisiae* however*, S. boulardii* has an optimal growth temperature of 37°C, increased acid tolerance and a robust cell wall that survives passage through the GI tract^18^. These physiological properties make *S. boulardii* well-suited for development as a synthetic biotic platform in humans.

Previous work has demonstrated the biosynthesis of small molecules (e.g., β-carotene, violacin)^9^, and recombinant proteins (e.g., HIV-GAG, lysozyme, IL-10, anti-*Clostridium difficile* nanobody tetramers) using engineered *S. boulardii* strains^11,19–24^. Currently, biopharmaceuticals or biologics, and in particular recombinant monoclonal antibodies (mAbs) or other antibody-like molecules, comprise more than 20% of new drug approvals and offer important new therapies for a variety of autoimmune diseases and cancers^25^. Notwithstanding these successes, antibody-based therapeutics present major challenges for therapeutic use including expensive manufacturing and quality control, product instability, requirements for a cold chain of storage and distribution, pharmacokinetic limitations, and injection-based delivery. These liabilities preclude access for many patient populations and impose substantial costs to healthcare systems.

Inflammatory bowel disease (IBD), including Crohn’s disease (CD) and ulcerative colitis (UC), are chronic inflammatory disorders of the GI tract caused by a combination of host genetics, environmental factors, and microbiome composition^26^. CD and UC are debilitating, life-long conditions that reduce quality of life and often result in complications that require surgery or other interventions. Tumor necrosis factor alpha (TNF, also known as TNFα) and other inflammatory cytokines are primary effectors of bowel inflammation and contribute to microbiome dysbiosis in CD^26^. A validated therapeutic approach is the administration of anti-TNF mAbs ^27^ to suppress the immune response. Although successful in achieving remission for some CD patients, treatment with an injectable mAb is cumbersome and expensive (> USD $10,000/patient/year)^28^. Moreover, mAbs against TNF have shortcomings that include failure to provide durable relief, unpredictable pharmacodynamics, systemic immune suppression, immunogenicity, and complex dosing regimens that preclude consistent attainment of therapeutically effective levels^29^. For these reasons, we focused on the development of *S. boulardii* as a synthetic biotic for the *in situ* production of neutralizing anti-TNF antibodies in the GI tract.

We optimized gene expression, protein secretion signals, and antibody-like scaffolds to build a robust *S. boulardii* synthetic biotic platform to produce biologics in the GI tract (**Fig. 1A**). Nanobodies (nAbs) based on a camelid heavy chain-single variable (VHH) scaffold yielded far higher levels of secretion that conventional antibody variants. We optimized the formulation and oral administration of *S. boulardii* for VHH delivery to the mouse GI tract. Administration of *S. boulardii* that secrete anti-TNF VHH to mice resulted in transient high-level colonization of the GI tract and *in situ* production of VHH at therapeutic concentrations without altering overall microbiome composition. An anti-TNF VHH-secreting strain significantly ameliorated disease symptoms and improved survival compared to control *S. boulardii* strains in a dextran sulfate sodium salt (DSS)-induced colitis mouse model. These results demonstrate localized production and delivery of a therapeutic VHH nAb via *S. boulardii* mitigates inflammatory disease activity in the gut.

**Figure 1.**
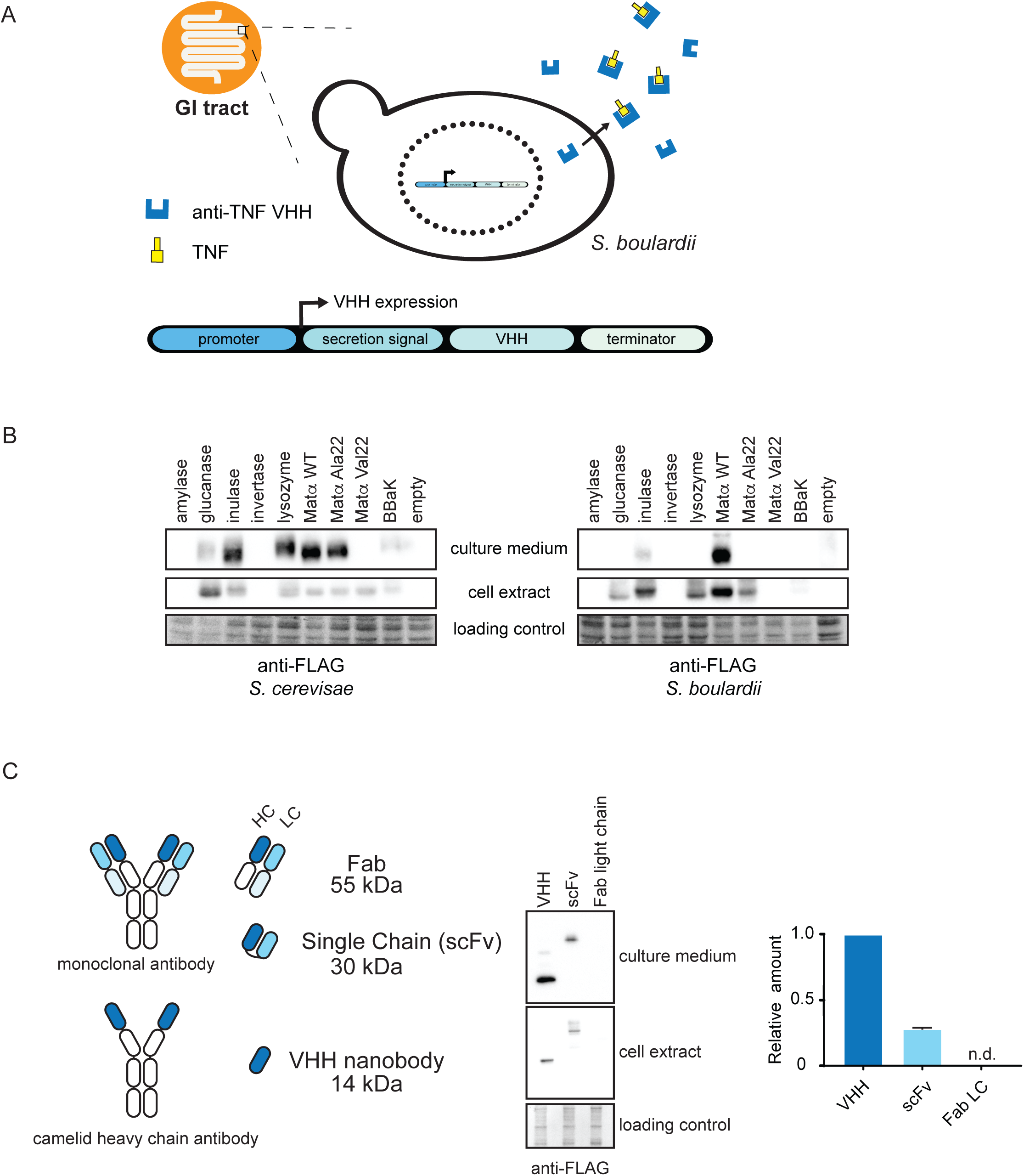
Optimization of secretion signals and antibody scaffold formats. **A)** Overview of *S. boulardii* synthetic biotic platform. **B)** Secretion signal efficiency. *S. cerevisiae* and *S. boulardii* were transformed with a galactose-inducible plasmid containing the light chain of a Fab’ against TcdA fused to nine secretion signals at the N-terminus and the FLAG epitope at the C-terminus. Cells were grown in SC-Ura media with 1% galactose + 1% raffinose at 30°C for 3 days in a shaker-reader. Secreted (culture medium) and retained (cell extract) Fab’ were detected by anti-FLAG immunoblot. Ponceau S stain was used as a loading control for cell extracts. **C)** S*. boulardii* were transformed with expression plasmids for a nanobody (VHH) against GFP, a single-chain antibody (scFv) against TcdA, or a light chain Fab’ fragment against TcdA, all fused to the Matα wild type secretion signal. Cells were grown in SC-Ura with 1% galactose + 1% raffinose at 30°C for 3 days. Retained and secreted Fab’ were detected by anti-FLAG immunoblot. Mean ± SD are indicated. n.d., not detected.

## Results

### Secretion tags and antibody scaffolds

Protein secretion requires the presence of a signal peptide at the N-terminus of the protein of interest to engage with the secretory pathway^30^. To enable the secretion of heterologous antibody scaffolds, a panel of nine candidate secretion signal peptide sequences was fused to the N-terminus of the light chain of a Fab’ dimer directed against *Clostridium difficile* toxin A (TcdA^31^) and tested for secretion yield. Secretion signals were derived from *S. cerevisiae* (amylase, glucanase, inulase, invertase), chicken (lysozyme), three variants of the *S. cerevisiae* Matα leader peptide (wild type; Matα variant App8 (‘Ala22a’) and variant Val22 (‘Val22a’)^32^, and a synthetic consensus signal peptide called BBaK^33^ (see **Supplementary Table 1** for all sequences and sources). These signal peptides were previously shown to support protein secretion of antibodies, antibody-like scaffolds, and other heterologous proteins in *S. cerevisiae*^34^. Signal sequence Fab’ constructs were expressed under the control of the *GAL1* inducible promoter on a 2 μm high copy *URA3* plasmid to alleviate potential toxic effects of Fab’ expression/secretion on yeast growth. As wild type *S. boulardii* yeast lacks the common auxotrophic markers available in *S. cerevisiae* laboratory strains, an auxotrophic *S. boulardii* strain was engineered by CRISPR-Cas9 mediated knockout of *URA3* to facilitate cloning and allow plasmid interoperability between *S. cerevisiae* and *S. boulardii*. To allow CRISPR-based gene engineering, we generated a plasmid that encodes *S. pyogenes* Cas9 and the *Aspergillus nidulans* acetamidase gene (*amdS*), a dominant prototrophic marker that allows *S. cerevisiae* to grow on acetamide as the sole nitrogen source^35^. After gene editing, removal of the Cas9 plasmid was achieved by counter selection on fluoroacetamide, which generates a toxic metabolite^35^. This method allowed scarless gene editing and subsequent elimination of the vector containing Cas9 and the sgRNA to minimize possible off-target cutting effects^36^. The signal peptide constructs were first tested using a Fab’ fragment in the wild type *S. cerevisiae* strain Sigma1278b. All signal peptides enabled secretion of the Fab’ fragment except for the amylase, invertase and MatαVal22 constructs (**Fig. 1B** left panel). The Matα wild type sequence was the most efficient secretion signal in *S. cerevisiae*, with only low amounts of detectable Fab’ fragments remaining inside the cell. In *S. boulardii*, only the Fab’ fragment fused to a Matα wild type signal peptide or, to a lesser extent, the inulase signal peptide, was expressed and secreted. We also observed higher retention of the Fab’ inside *S. boulardii* cells compared to the same constructs in *S. cerevisiae* cells (**Fig. 1B**, right panel). As the Matα leader sequence is conserved between *S. boulardii* and *S. cerevisiae*^14^ the basis for this difference is unclear. These results illustrate that even highly related yeast species can exhibit unexpected differences, underscoring the need for systematic characterization of chassis parts in biotechnological applications.

### Optimization of antibody scaffolds

Without considerable additional engineering of the secretion system, yeast species such as *S. cerevisiae* and *Pichia pastoris* are relatively inefficient in expressing canonical full-length antibodies^37^. We observed that while the Fab’ light chain readily produced detectable protein levels in *S. cerevisiae*, expression of a Fab’ heavy chain construct failed to produce detectable protein (data not shown). To overcome the requirement for Fab’ heterodimer expression, we assessed the production of human antibody-derived single-chain variable fragments (scFv) and monomeric antigen-binding VHH domains derived from cameloid species (sometimes referred to as nanobodies, nAbs). Both antibody-like scaffolds (**Fig. 1C**) have favorable properties compared to conventional antibodies or Fab fragments, including smaller size and improved protein stability while maintaining similar affinity for the target as conventional antibodies ^38^. To determine which antibody scaffold format was most compatible with *S. boulardii*, we compared the expression and secretion of a scFv against *Staphylococcal* enterotoxin B (scFv-GC132)^39^, a Fab against *C. difficile* toxin A (TcdA, ToxA-A03^31^) and a VHH against HIV-1 spike protein^40^. Scaffolds were fused to the Matα wild type secretion signal peptide and expressed under the control of *GAL1* promoter. In *S. boulardii*, the expression and secretion of the VHH construct was four times higher than the scFv construct, while the Fab light chain was not detectably expressed (**Fig. 1C**). The VHH format was therefore chosen as the preferred scaffold for further studies.

### Optimization of VHH production

Use of *S. boulardii* as a living cell microfactory platform to deliver protein-based therapeutic agents to the GI tract requires high level expression and secretion of the protein of interest. Promoters used to drive expression should ideally be active in the low oxygen and low glucose environment of the distal GI tract^41^. We considered the *ADH2* promoter (*ADH2^pr^*) as a potential constitutive promoter candidate as it is highly active at low glucose concentrations and under hypoxic conditions^42^. To assess the potential influence of protein sequence on expression, a panel of VHHs with different sequence characteristics was cloned into a 2 µm plasmid under the control of *ADH2^pr^*and N-terminally fused to the Matα wild type signal peptide^42^. For these experiments, we used the prototrophic *S. boulardii* parental strain (MYA-796) rather than a *ura3* auxotrophic derivative to maximize strain fitness. We expressed various *ADH2^pr^-VHH* constructs using a 2 µm plasmid bearing *kan* antibiotic selection that confers resistance to G418. This selection marker had the added benefit of allowing recovery and assessment of VHH-expressing *S. boulardii* strains after passage through the mouse GI tract (see below). *S. boulardii* strains bearing these plasmids expressed, secreted and properly processed the Matα-VHH fusion proteins at similar levels such that, on average, 95% of each VHH was secreted (**Fig. 2A**).

**Figure 2.**
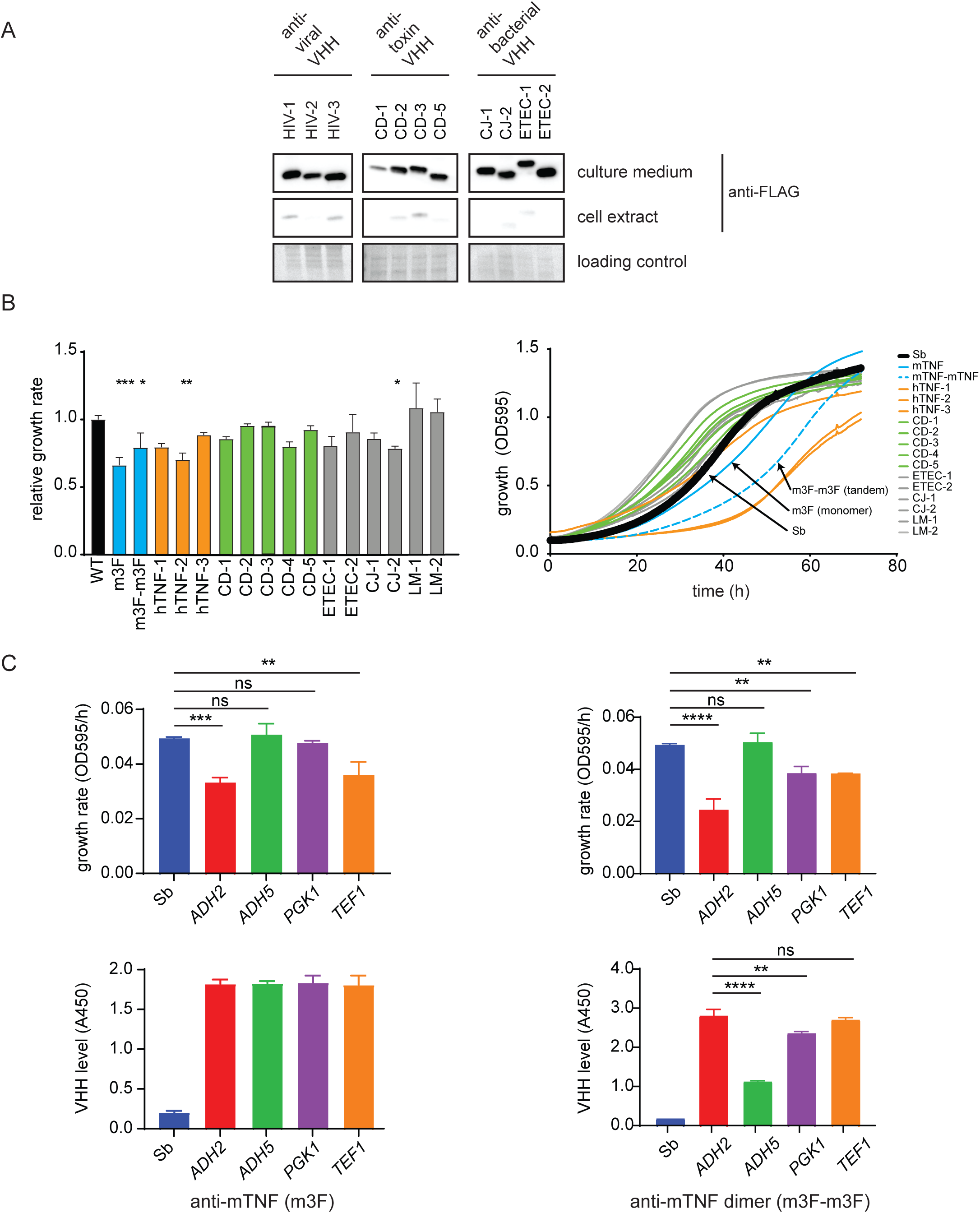
Characterization of secreted VHH nanobodies in *S. boulardii*. **A)** VHH nanobody secretion in *S. boulardii.* A panel of VHHs against HIV gp140 (HIV-1, HIV-2, HIV-3), *Clostridium difficile* TcdA (CD-1, CD-2, CD-3) and TcdB (CD-5) toxins and enterotoxigenic *E. coli* (ETEC) F4 fimbrae (CJ-1, CJ-2) and Stxe2 (ETEC-1, ETEC-2) were FLAG tagged and constitutively expressed from the ADH2 promoter in *S. boulardii* grown at 37°C for 3 days. Retained and secreted Fab’ from 60 µL of culture were detected by anti-FLAG immunoblot. Ponceau S stain was used as a loading control for cell extracts. **B)** Effect of VHH secretion on *S. boulardii* growth rate. A panel of the indicated VHHs directed against various antigens were N-terminally fused to the Matα wild type secretion signal and expressed from the *ADH2* promoter. Growth in liquid culture was monitored by OD_595_ and normalized to wild type growth rate based on curve slope during exponential phase. Statistical analysis was performed using Dunnett’s ANOVA test (* p<0.05, ** p<0.01, *** p<0.001). Mean ± SD are indicated. **C)** Optimization of anti-mTNF VHH expression in vitro. Anti-mTNF VHH (m3F) was expressed from the indicated yeast promoters. *Top*. Growth rates determined from growth curves. *Bottom*. Secreted mTNF VHH levels quantified by ELISA. Statistical analysis was performed using Tukey’s ANOVA test (* p<0.05, ** p<0.01, *** p<0.001, **** p <0.0001). Mean ± SD are indicated.

A reduction in fitness of engineered *S. boulardii* strains could cause a deleterious effect on growth *in vivo* and represent a potential liability for colonization of the GI tract, especially in competition with other *Saccharomyces* species in the host mycobiome^43^. To analyze the impact of VHH expression and secretion on *S. boulardii* fitness, we assessed yeast growth at human physiological temperature (37°C) and used growth rate as a proxy for fitness. Constitutive expression of VHHs did not impact *S. boulardii* growth rate in 12 out of 16 different VHH nanobodies tested (**Fig. 2B** and **Fig. S1**). Unfortunately, the VHHs that adversely affected fitness included an anti-mouse TNF VHH (m3F) and a corresponding tandem VHH dimer (m3F-m3F), which we had prioritized for use *in vivo* experiments (see below). For these constructs, we explored whether the promoter controlling VHH expression could have an impact on cell fitness. The growth rate of strains expressing either the m3F anti-mTNF VHH or the m3F-m3F VHH dimer, each under the control of four different promoters (*ADH2*, *ADH5*, *PGK1*, and *TEF1*)^42^, was compared during three days of growth at 37°C. ELISA was used to measure the amount of secreted VHH monomer or tandem VHH dimer after 3 days. For strains expressing the VHH monomer, the *ADH5* and *PGK1* promoters rescued full cell fitness without compromising VHH secretion. In contrast, although the use of the *ADH5* promoter rescued fitness for the VHH dimer strain, this was accompanied by a substantially decreased yield of secreted VHH dimer (**Fig. 2C**). These optimization steps yielded an *S. boulardii* strain that optimally expressed an anti-mTNF VHH monomer at 37°C without overt fitness defects.

### Optimization of VHH secretion

Deletion of the *HDA2*, *VPS5* and *TDA3* genes, which are implicated in trafficking and secretory functions, can increase protein secretion in *S. cerevisiae*^44^. The Cas9-*amdS* counter selection plasmid was used to generate CRISPR-mediated deletions for these three genes in *S. boulardii*. We analyzed the secretion of the anti-mTNF VHH under control of *ADH2*, *ADH5*, *PGK1*, *TEF1* and *TDH3* promoters. Secretion of each VHH was increased at 30°C in the individual knockout strains, except when expression was controlled by the *ADH2* promoter. At 37°C, the triple *hda2Δvps5Δtda3Δ* mutant expressing the anti-mTNF VHH under the control of *PGK1* promoter exhibited the highest level of VHH secretion, approximately 1.4-fold more than in the corresponding wild type strain (**Fig. S3**). However, all single, double and triple deletion mutants adversely affected cell growth rate at both 30°C and 37°C (**Fig. S3**). Due to this evident growth disadvantage compared to wild type *S. boulardii*, and the relatively minor improvement in VHH secretion yield, these gene deletions were not included in the final *S. boulardii* chassis used for VHH delivery *in vivo*.

### Secretion of functional anti-TNF VHH

We then quantified the yield of the secreted anti-TNF VHHs and tested for binding and neutralizing activity *in vitro*. We extended this analysis to human TNF (hTNF) in anticipation of future use of *S. boulardii*-based synthetic biotics in humans. VHHs directed against mTNF (i.e., m3F) and hTNF (termed hTNF−1, −2 and −3) were all produced and secreted (**Fig. 3A**). The maximum concentration in the culture supernatant was 6.9 μg/mL for the hTNF-1 VHH, a yield that is 6-fold higher than previously reported for an anti-hTNF VHH secreted by the Gram-positive bacterial probiotic *L. lactis*^4^. VHH oligomerization can increase apparent binding affinity to its target^38^, for example an anti-hTNF-3/anti-hTNF-1 VHH tandem dimer (called hTNF-4) has 1000 times more affinity for hTNF than the corresponding VHH domain alone^45^. Despite VHH dimer being similar in size to a scFv scaffold, *S. boulardii* secreted the VHH dimers at similar concentrations to the single VHHs, for both the hTNF-4 and m3F-m3F dimers (**Fig. 3A**). We used ELISA to demonstrate that the various secreted anti-TNF VHHs could bind recombinant TNF. The VHHs secreted by *S. boulardii* were functional and had a similar affinity (EC_50_) towards recombinant mouse and human TNF (**Fig. 3B**), as previously reported for *E. coli*-produced VHH^45,46^.

**Figure 3.**
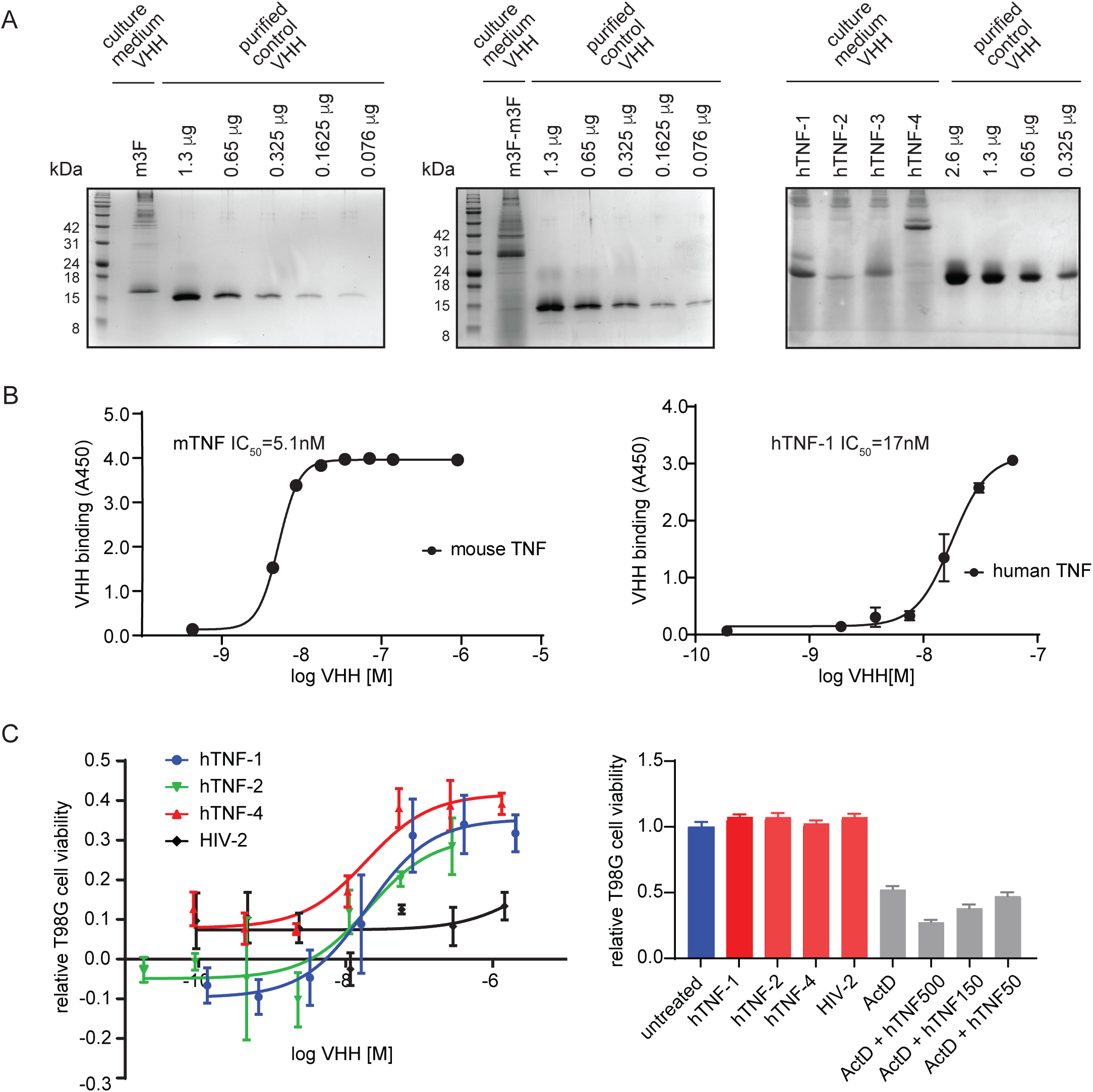
Characterization of anti-TNF VHH nanobodies produced in *S. boulardii*. **A)** Secreted levels of indicated VHH in yeast culture medium (60 µL) were quantified by comparison to a dilution series of purified recombinant His-tagged control nanobody (CD-2). Proteins were detected with Coomassie Brilliant Blue stain. *Left,* secreted anti-mTNF m3F VHH. *Middle, s*ecreted anti-mTNF m3F-m3F VHH tandem dimer. *Right*, secreted anti-hTNF VHH (hTNF-1, hTNF-2, hTNF-3) and anti-hTNF VHH tandem dimer (hTNF-4 = hTNF-3/hTNF-1). Estimated concentrations in culture medium were: 6.9 µg/mL for anti-mTNF m3F VHH; 4.4 µg/m for anti-mTNF m3F-m3F tandem dimer VHH; 5.5 µg/mL for hTNF-1 VHH; 1 µg/mL for hTNF-2 VHH; 3.4 µg/mL for hTNF-3 VHH; 4.9 µg/mL for hTNF-4 tandem VHH. **B)** Binding affinities of secreted anti-TNF VHH nanobodies. Binding of anti-mTNF VHH (m3F) to mTNF or anti-hTNF hTNF-1 VHH to hTNF was quantified by ELISA. Means ± SD are indicated. **C)** Neutralization of hTNF-induced cytotoxicity in human T98G glioblastoma cells treated with 40 nM actinomycin D to sensitize cells to hTNF. *Left*, effect of indicated anti-hTNH VHHs on viability of cells treated with 500 ng/mL hTNF. An anti-HIV-1 VHH (HIV-2) served as a non-specific VHH control. Mean ± SD are indicated. *Right*, cell viability 48 h after treatment with the highest concentrations of anti-hTNF VHHs used in neutralization assays (2.04 µM for hTNF-1; 0.28 µM for hTNF-2; 1.33 µM for hTNF-4; 1.45 µM for HIV-2). Viability effects of 40 nM actinomycin D alone or in the presence of the indicated concentrations of hTNF are also shown. Mean ± SD are indicated.

As a biological readout for the inhibition of hTNF by anti-hTNF VHH, we used a human T98G glioblastoma cell line assay in which actinomycin D was used to sensitize the cells to hTNF^47,48^. The anti-hTNF VHHs secreted by *S. boulardii* were able to protect against hTNF -induced cytotoxicity with high potency (EC_50_ < 100 nM) whereas an irrelevant control VHH had no effect (**Fig. 3C**, left). Importantly, we did not observe non-specific toxicity when T98G cells were treated with the VHHs alone (**Fig. 3C**, right). These results demonstrated that anti-hTNF VHHs secreted by *S. boulardii* retained neutralizing activity in vitro.

### S. boulardii colonization is influenced by the endogenous murine microbiome, pH and dose

Previous studies have suggested that *S. boulardii* does not stably colonize the murine GI tract in the absence of inflammation^17^. One possible explanation for this observation is that *S. boulardii* is partially outcompeted by the endogenous microbiome. To explore this possibility, C57BL/6 mice were treated with a single dose of streptomycin for 24 h, which is known to reduce colonization resistance of the mouse gut^49^. Mice were then gavaged with a single dose of 2×10^8^ colony forming units (CFUs) of *S. boulardii* resuspended in phosphate buffered saline (PBS; 10 mM phosphate pH 7.4, 137 mM NaCl, 2.7 mM KCl) and the residence time with or without streptomycin was determined by plating serial dilutions of resuspended feces on G418 medium to select for the *kan* resistance marker plasmid. In both streptomycin-pretreated and control mice, the *S. boulardii* CFUs recovered from feces peaked 24 h post-gavage and were undetectable after 72 h (**Fig. 4A**, left panel). However, when we examined the colonic tissue population of *S. boulardii* two days after gavage, streptomycin pretreated mice had almost two orders of magnitude higher *S. boulardii* colonization in the cecum and colon compared to the untreated controls (**Fig. 4A**, right panel).

**Figure 4.**
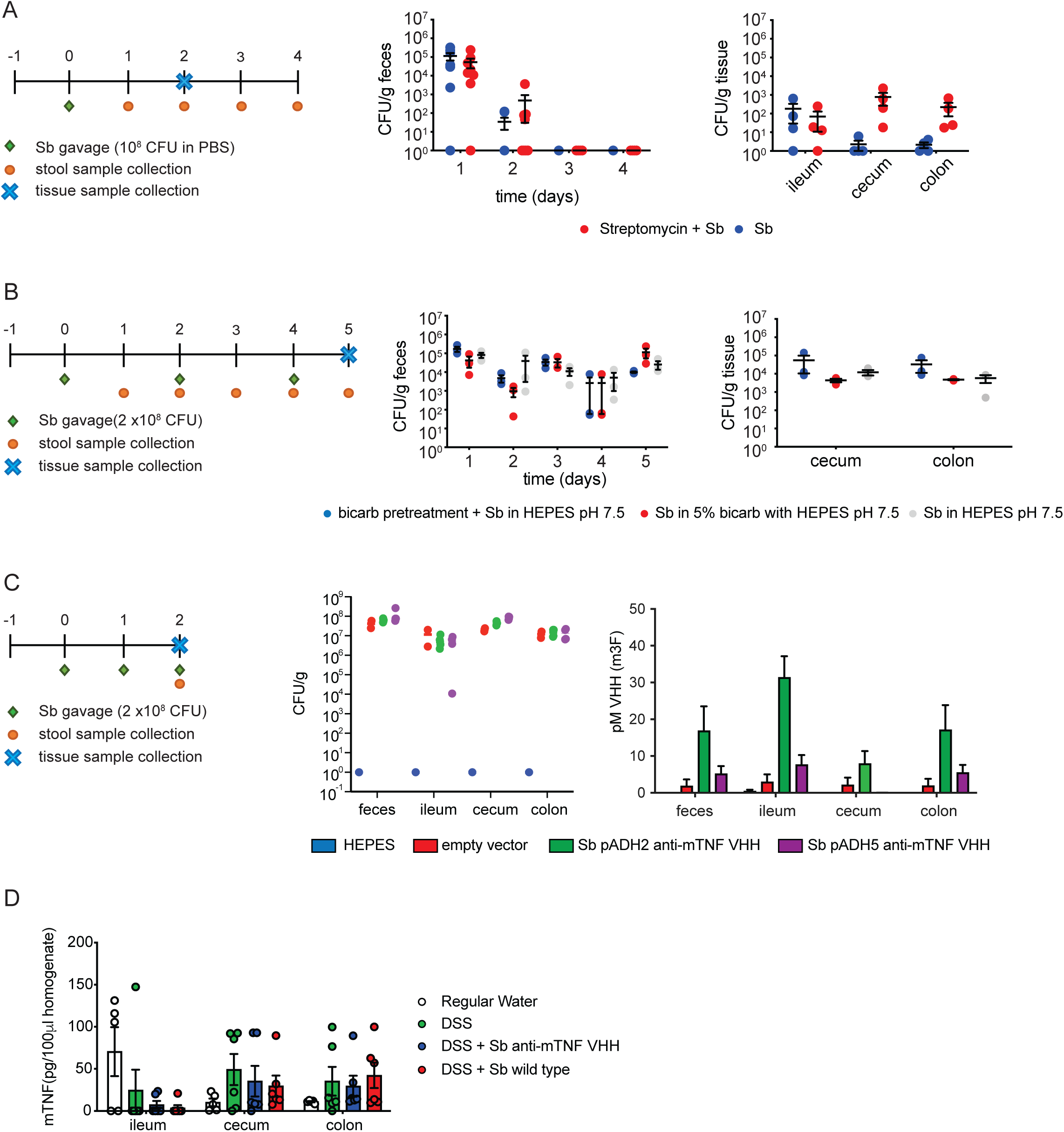
*In situ* production of anti-mTNF in the mouse GI tract by *S. boulardii*. **A)** Colonization of the GI tract by *S. boulardii*. Animals were untreated or pretreated with a single oral dose of 20 mg streptomycin 24 h prior to gavage with a single dose of 2 × 10^8^ CFU of *S. boulardii* in PBS bearing an empty vector. CFU were determined in feces by serial dilution for a 4 day time course and in the indicated tissues after 2 days. **B)** Effect of increased dose and pH neutralization on *S. boulardii* colonization. Animals were gavaged on days 0, 2 and 4 with 2×10^8^ CFU *S. boulardii* bearing an empty plasmid. Animals were either pretreated with 5% sodium bicarbonate prior to gavage in HEPES buffered saline (0.9% NaCl, 100 mM HEPES pH 7.5), or gavaged with 5% sodium bicarbonate + HEPES buffered saline, or gavaged with HEPES buffered saline only. CFUs were determined by serial dilution. Mean ± SD are indicated. **C)** Quantification of anti-mTNF VHH production from two different promoters in the GI tract. Animals were pretreated with a single oral dose of 20 mg streptomycin 24 h prior to gavage. Animals were gavaged on days 0, 1 and 2 with 2 × 10^8^ CFU *S. boulardii* transformed with an empty plasmid or plasmids expressing anti-mTNF VHH under the control of the *ADH2* or *ADH5* promoter. *Left, S. boulardii* CFU per gram of feces or tissue after 3 days of treatment under the HEPES buffered saline regime for indicated strains. *Right,* anti-mTNF VHH quantification in feces and tissues by ELISA for indicated strains after three days of treatment. Purified anti-mTNF VHH (m3F) produced in *E.coli* was used to generate a standard curve for ELISA. **D)** Quantification of mTNF levels in tissues by ELISA.

Despite better survival of *S. boulardii* over *S. cerevisiae* in acidic conditions^17^, the low pH of the stomach might still affect *S. boulardii* fitness and decrease colonization rates. To test whether delivery in more alkaline conditions might improve colonization, mice were pretreated with a 5% bicarbonate buffered solution followed by gavage with wild type *S. boulardii* in HEPES buffered saline (0.9% saline, 100 mM HEPES pH 7.5), or with Sb in 5% bicarbonate and HEPES buffer, or with Sb in HEPES buffered saline only. To further increase colonization, we also doubled the gavage dose of *S. boulardii* to 2×10^8^ CFU and repeated the gavage every other day for 5 days. Higher levels of colonization in tissue were observed with each of these pH neutralization schemes (**Fig. 4B**). Compared to the lower dose of *S. boulardii* in PBS, the higher dose combined with pH neutralization increased colonization 30-fold (from 7×10^2^ to 2.3×10^4^ CFU/g tissue) in the cecum and 70-fold (from 2×10^2^ to 1.4×10^4^ CFU/g tissue) in the colon (**Fig. 4A** versus **B**). For simplicity of administration, subsequent experiments used HEPES buffered saline alone.

### In vivo synthesis of VHH nanobodies by S. boulardii

We observed above that *S. boulardii* fitness and VHH secretion were impacted by the choice of promoter. To explore if these *in vitro* results extended to GI tract colonization *in vivo*, *S. boulardi*i expressing anti-mTNF VHH under the control of the *ADH2* or *ADH5* promoter, or *S. boulardii* with an empty plasmid vector, or HEPES buffered saline alone were gavaged for 3 successive days. Three days after the first gavage, VHH content was analyzed in the ileum, cecum, colon and feces by ELISA. *S. boulardii* that expressed anti-mTNF VHH from the *ADH2* promoter yielded higher VHH levels in all tissues analyzed, with a 3-fold increase in the colon compared to expression from the *ADH5* promoter (245± 96.86 pg/mL for *ADH2* versus 79 ± 30.48 pg/mL for *ADH5*) (**Fig 4C**, right panel). This increased level of secretion was not due to differential colonization of the GI tract between the two strains (**Fig. 4C**, left panel). We therefore prioritized the *ADH2*-driven anti-mTNF VHH expression strains for subsequent testing in a mouse model of colitis.

A critical question for *in vivo* efficacy was whether sufficient levels of neutralizing VHH are produced compared to mTNF. We therefore quantified relative anti-mTNF VHH and mTNF levels by ELISA and converted these values to concentration estimates using standard curves with recombinant proteins (see Methods). Because mTNF levels in the uninflamed colon were expected to be low, we assessed mTNF levels in DSS-treated animals, which exhibit a high degree of colonic inflammation. We found that anti-mTNF VHH levels in tissues from untreated animals, as driven by the *ADH2* promoter, exceeded the levels of mTNF (**Fig. 4C** versus **4D**; between 25-50 pg/100 uL TNF detected by ELISA in feces or different tissues = 5-10 pM of active TNF trimer, compared to 10-30 pM of anti-TNF VHH detected in feces or different tissues). These results suggested that strains producing anti-mTNF VHH would have the potential to neutralize endogenous mTNF and remediate GI inflammation in the DDS model.

### Passage of S. boulardii through the intestinal tract does not affect VHH production or stability

An important question for any synthetic biotic is whether the intestinal environment selects against the production of the desired biologically active agent. To address this issue, we compared anti-mTNF VHH levels produced by input *S. boulardii* strains to strains isolated as individual clones from feces 2 days post-gavage. We found no difference in VHH production between the input and output strains recovered after passage through the GI tract (**Fig. S2A**). This result suggested that there was negligible counter-selective pressure against VHH-producing *S. boulardii* in the GI tract. A further question was whether the secreted VHH was stable in the intestinal environment. To address this question, we assessed the stability of the anti-mTNF VHH when spiked into GI extracts. The anti-mTNF VHH was stable in feces, and in cecum and colon extracts, but was degraded when incubated with an ileum extract, which is estimated to have 20-fold more proteolytic activity than the feces^50^ (**Fig S2B**). We concluded that the activity of the *S. boulardii*-produced anti-mTNF VHH would likely not be impaired in the large intestine.

### DSS-induced colitis does alter S. boulardii colonization or anti-mTNF VHH production

To assess efficacy *in vivo*, we evaluated the anti-mTNF VHH strain in the well-established dextran sulfate sodium (DSS)-induced mouse colitis model, which is commonly used to evaluate anti-inflammatory therapeutic strategies in IBD^4,5,51^. DSS is a sulfated polysaccharide that is toxic to colonic epithelium and when administered to mice in drinking water causes disruption of the intestinal epithelial barrier and inflammation limited to the colon^52^. We first determined whether DSS treatment altered *S. boulardii* colonization of the GI tract. We gavaged C57BL/6 mice once every two days with 2×10^8^ CFU of the anti-mTNF strain in HEPES buffered saline for 7 days in the presence or absence of 3% DSS in drinking water (**Fig. 5A**). DSS did not cause any significant difference in CFU counts in either feces or colonic tissue (**Fig. 5B**). Although it has been previously shown that *S. boulardii* retention in the mouse intestinal epithelium is increased by inflammation^17^, we did not observe any significant increase in *S. boulardii* CFUs in cecum and colon tissue upon DSS-induced inflammation. We then determined VHH levels in tissues at experimental endpoint of the DSS treatment time course. As controls, we used a strain that secreted an irrelevant VHH (anti-Gp140 HIV^40^, termed HIV-2 above) or treatment with HEPES buffered saline only. The secreted anti-mTNF VHH and control VHH were readily detected by ELISA in feces, as well as in in cecum and colonic fractions (**Fig. 5C**). This result demonstrated that VHH secretion *in vivo* was not affected by DSS-induced colitis.

**Figure 5.**
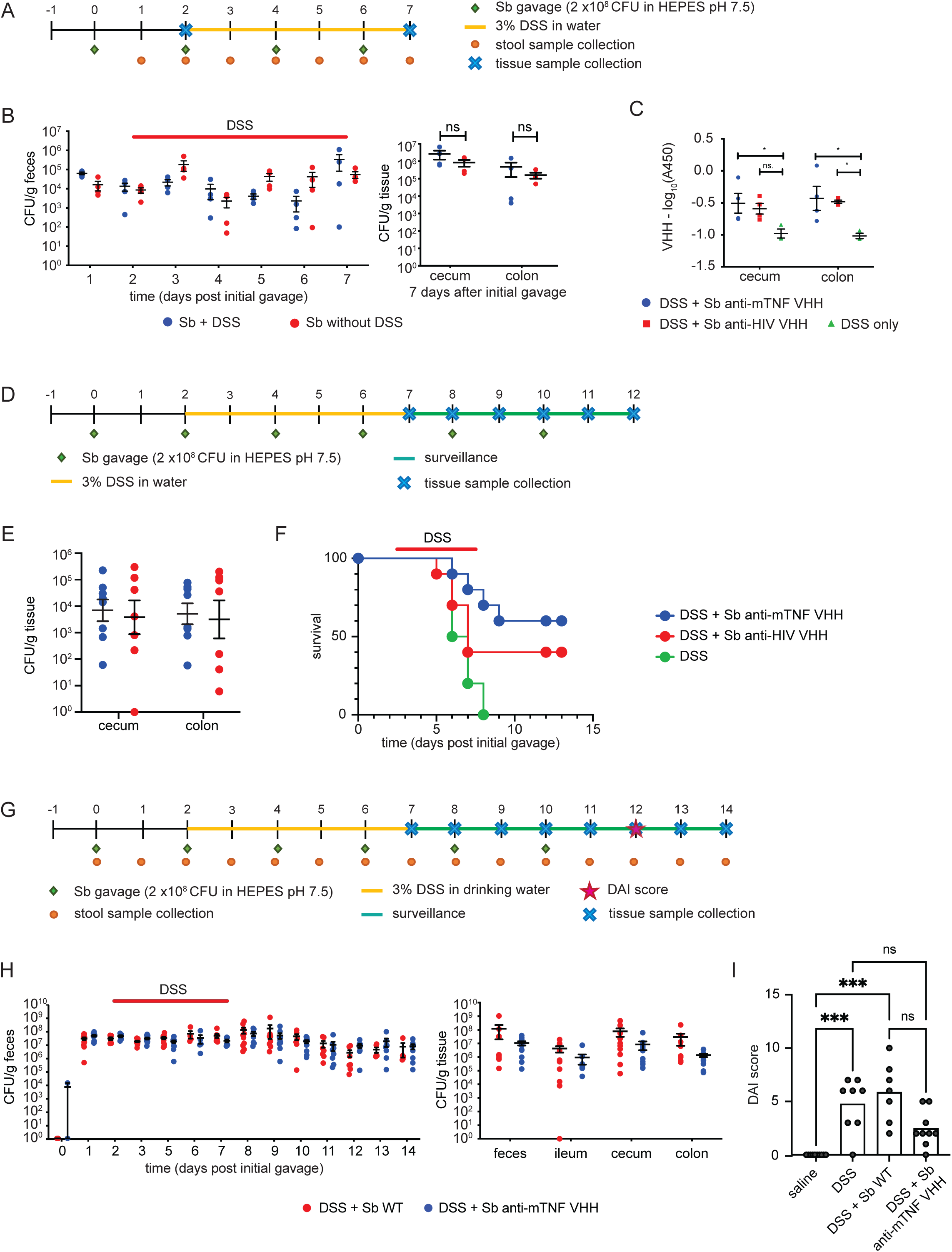
*In situ* production of anti-mTNF VHH by *S. boulardii* increases survival and improves symptoms in a DSS colitis model. **A)** Schematic of experimental time course for *S. boulardii* colonization and VHH production. **B**) *S. boulardii* colonization with or without DSS-induced colitis in feces over 7 day time course (*left*) and in GI tissue (*right*) at 7-day experimental endpoint. Animals were pretreated with a single dose of 20 mg streptomycin 24 h prior to initial *S. boulardii* administration then gavaged once every 2 days with 2×10^8^ CFU wild type *S. boulardii* or *S. boulardii* expressing anti-mTNF VHH from the *ADH2* promoter in HEPES buffered saline. CFUs were determined by serial dilution of feces at each time point or in indicated tissues at either humane (>20% weight loss) or experimental endpoint. Means ± SEM are shown. **C)** Relative levels of anti-mTNF VHH or control anti-HIV-1 VHH (HIV-2) in tissues at 7-day experimental endpoint as determined by ELISA. **D)** Schematic of 12 day experimental time course for survival after DSS treatment. **E)** *S. boulardii* colonization with or without DSS-induced colitis in feces GI tissue at 12-day experimental endpoint. Animals were pretreated with a single dose of 20 mg streptomycin 24 h prior to initial *S. boulardii* administration then gavaged once every 2 days with 10^8^ CFU *S. boulardii* expressing anti-mTNF VHH (m3F) or control anti-HIV-1 VHH (HIV-2) from the *ADH2* promoter in HEPES buffered saline. CFUs were determined in tissues at either humane (>20% weight loss) or experimental endpoint. Each datapoint indicates an individual animal. Means ± SEM are shown. **F).** Survival curves after DDS-induced colitis with administration of *S. boulardii* secreting either anti-mTNF VHH or control anti-HIV VHH versus HEPES buffered saline alone. **G)** Schematic of 14 day experimental time course for disease severity. Animals were pretreated with a single dose of 20 mg streptomycin 24 h prior to initial *S. boulardii* administration then gavaged once every 2 days with 10^8^ CFU wild type *S. boulardii* or *S. boulardii* expressing anti-mTNF VHH (m3F) from the *ADH2* promoter in HEPES buffered saline. CFU were determined by serial dilution of feces at each time point or in indicated tissues at either humane (>20% weight loss) or experimental endpoint. **H)** *S. boulardii* colonization with or without DSS-induced colitis in feces (*left*) and GI tissue (*right*) at 14 day experimental endpoint. Means ± SEM are shown. **I)** Disease activity index (DAI) scores after DSS-induced colitis with administration of *S. boulardii* secreting either anti-mTNF VHH (m3F) or control anti-HIV VHH (HIV-2) versus HEPES buffered saline alone or HEPES buffered saline without DSS treatment. Animals were pretreated with a single dose of 20 mg streptomycin 24 h prior to the first *S. boulardii* administration. DAI was scored at day 12 of the time course. DAI scores were: DSS + vehicle only, 5.43; wild type *S. boulardii*, 5.86; *S. boulardii* anti-mTNF VHH (m3F) 2.44; water, 0. Statistical analysis was by Mann-Whitney T-test (p < 0.05, ** p < 0.01, *** p<0.001, **** p<0.0001). Means ± SEM are indicated.

### S. boulardii expressing anti-mTNF VHH reduces disease severity in DSS-induced colitis

To assess whether in situ secretion of anti-mTNF reduced mortality in the DDS model, we carried out a 12 day time course experiment in which animals were allowed to progress after DSS treatment (**Fig. 5D**). Animals were gavaged once every 2 days with 2×10^8^ CFU of *S. boulardii* in HEPES buffered saline for 7 days in the presence of 3% DSS in drinking water from days 2 to 7, followed by 5 days of daily gavage in the absence of DSS. Animals treated with a strain expressing the anti-HIV VHH or with HEPES buffered saline only were used as controls. No difference in *S. boulardii* retention was detected in GI tissues at the end of the experiment (**Fig. 5E**). Administration of the anti-mTNF VHH strain increased survival of mice treated with DSS by 60% compared to administration of HEPES buffered saline alone (Gehan-Breslow-Wilcoxon test, p = 0.0022) (**Fig. 5F**). The control VHH strain conferred a survival benefit of 40% over HEPES buffered saline alone but this effect was lower than for the anti-mTNF VHH strain (control VHH versus anti-mTNF VHH, p= 0.0087, log-rank Mantel-Cox test). To further characterize the effect of the anti-mTNF VHH strain, we performed a second time course-recovery experiment for 14 days and monitored disease activity index (DAI) with a strain bearing an empty vector as the control (**Fig. 5G**). As expected, no differences in colonization between the strains were observed in feces or tissues (**Fig. 5H**). However, as measured by DAI score, the anti-mTNF VHH strain had a significant benefit over both the vector only control strain and the HEPES buffered saline control in reducing disease severity at the end of the experiment (anti-mTNF versus empty vector strain p=0.0001; anti-mTNF versus HEPES only p=0.0104) (**Fig. 5I**). Collectively, these results demonstrated that in situ production of the anti-mTNF VHH by *S. boulardii* had a pronounced beneficial effect on both survival and disease severity in the DSS model.

### S. boulardii expressing anti-mTNF VHH does not alter the endogenous microbiome

Long term administration *S. boulardii* in the mouse DSS colitis model has been previously shown to increase bacterial diversity of the microbiome^53^. We thus explored whether mice treated with *S. boulardii* expressing anti-mTNF VHH exhibited alterations in the microbiome. Mice were treated with strains expressing anti-mTNF VHH or control anti-HIV^Gp140^ VHH, or HEPES buffered saline alone, administered DSS for 5 days (**Fig. 6A**), and then evaluated for alterations to the microbiome composition by 16S rRNA amplicon sequencing. As assessed by Shannon diversity index, which measures alpha diversity of the intestinal microbiome, we observed no significant difference between mice administered the buffer control versus strains expressing either VHH (**Fig. 6B**). Thus, the engineered probiotics did not influence the diversity of the intestinal microbiota. The similarity between microbiome composition (i.e., beta-diversity) of the different sample groups was then assessed using canonical correspondence analysis (CCA). Dispersion of beta diversity was computed using permutational multivariate analysis of variance (PERMANOVA). We observed that samples taken prior to colitis induction, after DSS treatment or at endpoint clustered together independently of the treatment (**Fig. 6C**). Based on these results, we concluded that the beneficial effects of acute treatment with the anti-mTNF VHH strain were due to the anti-mTNF VHH itself and not a general effect of *S. boulardii* on the microbiome. We also concluded that short term administration of the anti-mTNF VHH strain did not disrupt the overall composition of the microbiome.

**Figure 6.**
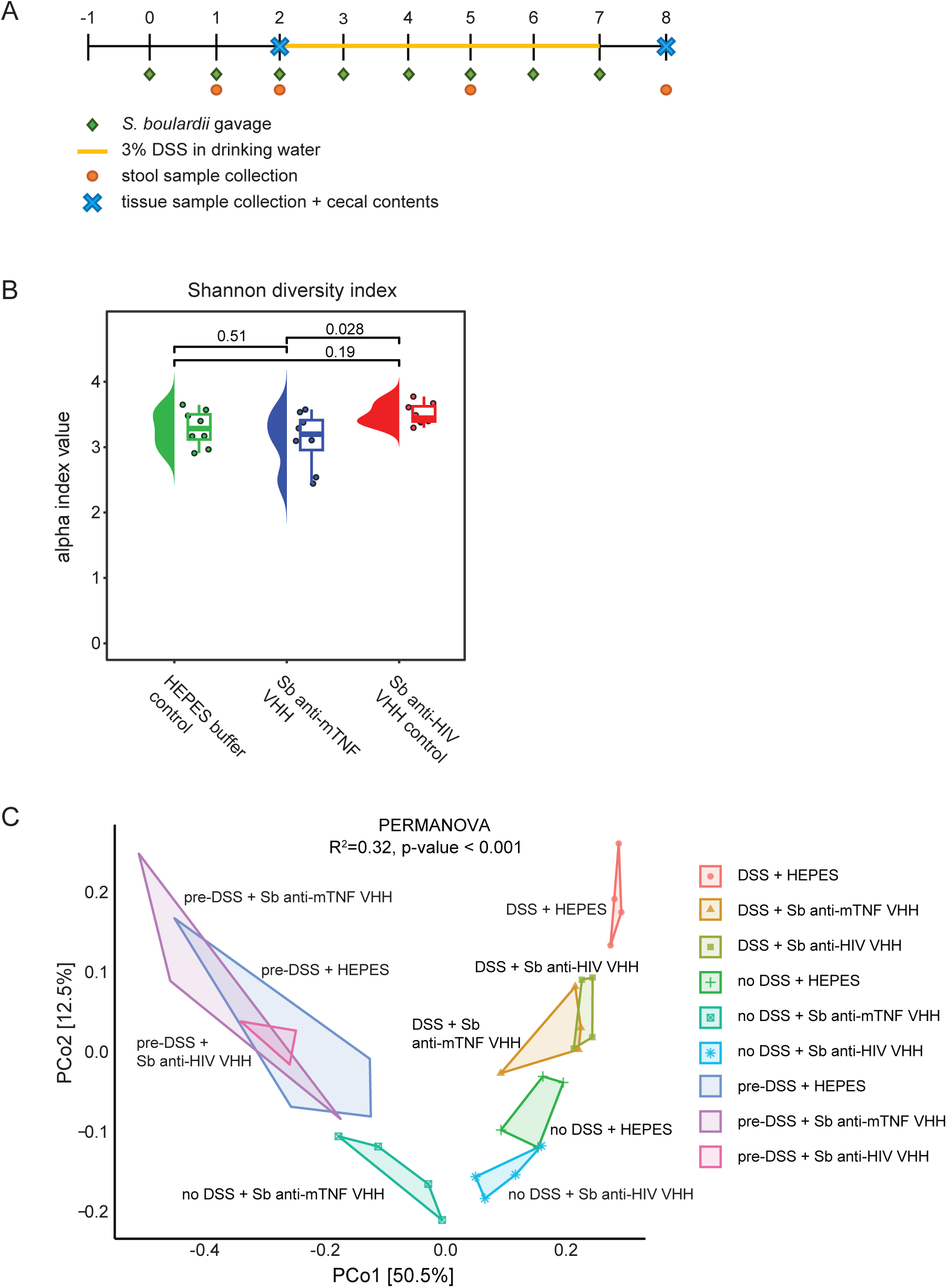
Assessment of the endogenous GI microbiome in the presence or absence of *S. boulardii* expressing anti-mTNF VHH. Cecal contents from animals administered the indicated *S. boulardii strains* with or without DSS-induced colitis were sequenced in the V3-4 region of bacterial 16S rRNA. **A)** Schematic of 8 day experimental time course. **B)** Alpha diversity assessed by Shannon index. Statistical significance was assessed by ANOVA. **C)** Beta diversity assessed by principal component analysis of amplicon sequence variants. Statistical significance was assessed by PERMANOVA.

## Discussion

Wild type *S. boulardii* has been extensively studied and used as a probiotic for the treatment of antibiotic-induced diarrhea, *Clostridium difficile* infections, as well as in patients suffering from ulcerative colitis and Crohn’s disease^15,54^. The engineering of *S. boulardii* to deliver therapeutic biologics such as nanobodies directly to the GI tract has the potential to combine the general therapeutic effect of this commonly used yeast probiotic with highly targeted therapies^2,23^. Importantly, the use of *S. boulardii* as a living cell microfactory for *in situ* production of VHH nanobodies, may provide an accessible and cost-effective alternative to conventional recombinant biologic therapies that currently require systemic intravenous administration. Here, we demonstrate that anti-mTNF VHH nanobodies secreted in the GI tract can alleviate symptoms in a mouse model of colitis. Our platform was optimized for (i) VHH nanobody expression under low nutrient conditions that mimic the GI environment, (ii) maximal levels of VHH nanobody secretion without causing proteostatic stress, and (iii) delivery regimes that maximize transient population of the GI tract. We specifically avoided use of auxotrophic markers for strain engineering and instead used a fully prototrophic wild type strain to maximize strain fitness in the GI environment, as enabled by a combination of CRISPR/Cas9-based genome engineering and the dominant counter-selectable *amdS* marker^55^. These features represent improvements over previous engineering methods in *S. boulardii* based on laterally transferrable plasmid vectors and dominant antibiotic resistance markers^21,56^. Our results also underscore the importance of optimization for *in vitro* versus *in vivo* applications. Notably, while the *ADH2* and *ADH5* promoters yielded similar amounts of VHH in culture media, the *ADH2* promoter clearly outperformed *ADH5* for the production of VHH in tissues of the GI tract, despite no difference in CFU levels between the two strains.

Tandem bivalent or multivalent versions of VHH nanobodies can improve the affinity toward target epitopes^12^. For example, high avidity nanobody multimers can neutralize influenza, SARS-CoV-2 and *C. difficile*^12,57,58^. We tested several anti-human and anti-mouse TNF bivalent tandem VHH constructs but found that secretion of larger fusion proteins tended to reduce *S. boulardii* fitness, likely due to proteotoxic stress. While secretion of bivalent VHHs might be improved by engineering the *S. boulardii* protein secretion system^37,44^, modifications to *S. boulardii* need to be carefully assessed for impact on both secretion efficiency and fitness. Despite the close relationship between *S. boulardii* and *S. cerevisiae*, our secretion signal optimization experiments demonstrated that improvements in *S. cerevisiae* do not necessarily translate to *S. boulardii*, as shown by the deleterious effects of deleting *HDA2*, *VPS5* and *TDA3* in *S. boulardii.* These unexpected negative effects may partly be because the optimization of *S. cerevisiae* protein secretion has been in the context of improving yields of recombinant proteins under optimal growth conditions in fermenters. In the context of synthetic biotic development, *S. boulardii* secretion needs to be compatible with the relatively nutrient-poor conditions in the GI tract and competitive fitness with the complex GI microbiome.

*S. boulardii* is a generally recognized as safe agent and we observed no adverse effects upon oral administration of either wild type or engineered *S. boulardii* to mice. In addition, we did not observe any significant changes in the microbiome associated with the acute administration of either wild type or engineered *S. boulardii* strains. Further advantages of *S. boulardii* as a synthetic biotic platform include its relatively short residency time in the GI tract and the absence of horizontal gene transfer that is a risk with bacterial synthetic biotics^59^. Our results demonstrate the pre-clinical efficacy of engineered *S. boulardii* as a biotherapeutic production platform for anti-TNF VHH-based remediation of IBD, positioning this approach for consideration in future human clinical trials.

The *S. boulardii* platform developed here can be readily adapted for the delivery of other therapeutic biologics. For example, *S. boulardii*-mediated secretion of neutralizing nanobodies directed at targets implicated in IBD such as IL-12/IL-23 and α4β7/α4β1 integrins may have advantages compared to systemic delivery of approved monoclonal antibodies against these targets^60^. This approach may be extended to *S. boulardii*-mediated delivery of nanobodies that neutralize bacterial antigens associated with IBD pathogenesis^61^. The *S. boulardii* platform may also be readily deployed for veterinary applications, for instance, in the prevention and treatment of various diseases and infections in livestock. *S. boulardii* probiotics are already approved for use in livestock to increase production, reduce cost, and mitigate dependence on antibiotics^62–66^. Further development of the *S. boulardii* platform holds promise for the low-cost production and facile oral delivery of highly specific biologic agents in manifold contexts.

## Methods

### Plasmid construction

All constructs consisted of a promoter, secretion signal, antibody, secretion tag, *CYC1* terminator, selection marker, 2 µm origin of replication, ampicillin resistance gene and *E. coli* origin (see **Fig. S4A** for plasmid map). Promoters were amplified from *S. cerevisiae* or *S. boulardii* and secretion signals (Supplementary Table 1) and nanobodies (Supplementary Table 2) were obtained as gBlocks (Integrated DNA Technologies). Antibody sequences were as reported^31,39^. Plasmids were constructed using Gibson Assembly cloning^67^. To generate gene knockouts, the plasmid pGZ110 that expresses SpCas9 and a targeting sgRNA (gift from Bruce Futcher, SUNY Stony Brook) was modified to replace the *URA3* auxotrophic selection marker with the *amdS* recyclable selectable marker, which confers resistance to acetamide and can be counter-selected using fluoroacetamide^35^. Guides to target *URA3*, *HDA2*, *TDA3* and *VPS5* were designed using the Synthego Knockout Guide Design tool (https://design.synthego.com/#/). The vector also contained a homology repair region that introduced several stop codons and a frameshift to prevent readthrough into each coding region, as well as a PAM site deletion to prevent continuous Cas9 nucleolytic activity (see Supplementary Table 3 for sequences). After homozygotic deletion was confirmed by Sanger sequencing (**Fig. S3A)**, the plasmid was removed by counter-selection on fluoroacetamide medium. All plasmids used in this work are listed in Supplementary Table 4.

### Yeast strain construction

Yeast cells were transformed using the lithium acetate method^68^. *S. boulardii* (MYA-796, obtained from ATCC) and *S. cerevisiae* (Sigma1278b) were inoculated overnight at 30°C with shaking at 220 rpm, then diluted to an OD_595_ of 0.5, and incubated at 30°C and 220 rpm until an OD_595_ of 1.5 was reached. Cells were then washed with distilled water, resuspended in transformation mix (240 µL 50% PEG 3350, 36 µL 1M lithium acetate, 50 µL single-stranded carrier DNA (2 mg/mL in TE), 36 µL 1M DTT, and 16 µL water + plasmid DNA (1 μg in *S. boulardii* and 100 ng in *S. cerevisiae*). Cells were incubated at 42°C for 40 min, pelleted and washed with 1 mL sterile water, re-suspended in 200 µL water and immediately plated onto appropriate selective medium. For G418 selection cells were recovered overnight in XY rich medium (YEPD + 0.01% w/v adenine and 0.02% w/v tryptophan) at 30°C and 220 rpm. The transformation efficiency of *S. boulardii* was typically 30 to 50-fold lower than for *S. cerevisiae*^36^. All strains used in this work are listed in Supplementary Table 5.

### Antibody and nanobody detection

Cells expressing antibody fragments or VHH nanobodies under the control of the *GAL1* promoter were precultured overnight in XY + 2% raffinose at 30°C, diluted 200-fold into XY + 2% galactose for 2-4 days. Cells expressing antibodies/nanobodies under the control of constitutive promoters were grown for 3 days in XY + 2% glycerol+ 2% ethanol or XY + glucose at the indicated concentrations and at the indicated temperatures. Cultures were centrifuged, supernatant collected, and pellets washed with cold water. Proteins in the culture medium were precipitated in 10% TCA dissolved in acetone at a 1:10 ratio. Cell pellets were dissolved with 2 M LiOAc and 0.4 M NaOH for five min in each solution^69^. For immunoblot analysis, antibodies/nanobodies were resolved by SDS-PAGE containing 15% acrylamide (VHH) or 12% acrylamide (Fab’) (BioRad) then transferred to PVDF membrane using a wet transfer system (Bio Rad) or directly stained with Coomassie Blue. Membranes were probed with mouse monoclonal anti-FLAG-HRP (Sigma A8592) at 1:5000 dilution or anti-VHH-HRP (GenScript A01861) at 1:3000 dilution and signals visualized by chemiluminescence (Western Lightning Plus-ECL) and a ChemiDoc imaging system (BioRad).

### Growth rate analysis

A preculture was grown overnight until saturation in XY + 2% ethanol+ 2%glycerol or 2% glucose then diluted 20-fold in 96-well clear flat-bottomed microplates (Corning Costar) in the same medium. The plates were sealed and incubated for 72 h on a Sunrise shaker-reader (Tecan) at 37°C with shaking at 564 rpm. Absorbance at 595 nm was measured every 15 min for 24 h. Growth rate was calculated as the linear portion of the curve using PRISM software.

### VHH nanobody purification

Cells were grown in XY + 2% ethanol + 2% glycerol for 3 days in 500 mL culture at 30°C. The culture was centrifuged at 3000 rpm and the supernatant was adjusted to pH 8.0 with 10 N NaOH. The supernatant was applied on a His-Trap HP column (GE Healthcare), washed with buffer (50 mM Tris pH 8.0, 120 mM NaCl, 10% glycerol, 20 mM imidazole), and eluted with wash buffer with 500 mM imidazole. Eluted fractions were concentrated by ultrafiltration using a 3 kDa cutoff spin filter (Vivaspin). Quantification of VHH abundance was performed using a Pierce 660 nm colorimetric assay (ThermoFisher).

### Direct ELISA

For proteins in yeast culture medium, 100 ng of either mTNF or mouse monoclonal anti-VHH in PBS were incubated for 2 h at 4°C on protein-binding Nunc MaxiSorp® flat-bottom 96-well plates and then blocked with 100 µL of 0.1 M NaHCO_3_ pH 8.6 + 2% BSA for 1 h at room temperature (RT). The protein of interest was diluted in PBS and incubated with the plate for 1 h at RT, followed by incubation with anti-FLAG-HRP or anti-HIS-HRP (1:5000 in PBS) for 1 h at RT and subsequent development with 50 µL of 3,3′,5,5′-tetramethylbenzidine (TMB, R&D Systems #555214) substrate. The reaction was stopped with 100 µL of 1 M sulfuric acid and the plate was read in a microplate reader (Tecan). Absorbance at 450 nm on an EnVision microplate reader (Perkin Elmer) was used to determine the amount of bound antibody.

For mouse tissues, 100 ng of the protein of interest in PBS was coated overnight at 4°C on protein-binding Nunc MaxiSorp® flat-bottom 96-well plates and then blocked with 100 µL of FBS blocking ELISA/ELISPOT diluent (eBioscience). To generate tissue samples, female C57BL/6N (Charles River) 6- to 8-week-old mice were gavaged with 20 mg streptomycin then administered ∼10^8^ CFU daily doses of saturated *S. boulardii* yeast cultures via oral gavage 1 day and 3 days later. Mice were euthanized 1 day later, and the ileum, cecum and colon tissues were removed. Fecal pellets and tissues, including luminal contents, were homogenized in 1 mL PBS using a Retsch mixer mill homogenizer (RM400). Samples were centrifuged and the amount of mF3 VHH nanobody in the supernatants was quantified using a standard curve generated with purified m3F-FLAG VHH. For ELISA, an anti-FLAG capture antibody (Genescript #A00170) was used at 1:4000 dilution, followed by an anti-VHH-HRP detection antibody (Genescript # A01861) at 1:4000. ELISAs were developed using TMB (R&D Systems #555214) and read at 450 and 570 nm using an Envision plate reader (Perkin Elmer).

### Nanobody toxicity assays

T98G cells (ATCC) were grown in Dulbecco’s modified Eagle’s medium supplemented with 100 units/mL penicillin, 100 μg/mL streptomycin and 10% heat-inactivated fetal bovine serum. Cells were maintained at 37°C in 5% CO_2_ and 100% humidity. For cytotoxicity assays, cells were seeded in 384-well plates for 24 h, followed by VHH addition using an Echo 555 acoustic dispenser (Labcyte). Cell viability was evaluated after 48 h by Cell Titer Glo in a Tecan reader (luminescence, gain 135, integration time 40 s). To evaluate the neutralization potential of different VHH preparations, a combination of actinomycin D (40 nM) and hTNF (500 ng/mL) was empirically chosen to provide an optimal dynamic range for the assay. Cells were treated with 500 ng/mL hTNF and 40 nM actinomycin D in the presence of increasing concentrations of the VHH purified from yeast culture medium. Neutralization activity was calculated by normalizing cell viability in actinomycin D as 1 and cell viability in actinomycin D + hTNF as 0.

### Colonization of mice with S. boulardii

All mouse experiments were performed in the Central Animal Facility at McMaster University under animal use protocol 20-12-41 as approved by the Animal Research Ethics Board and under compliance with Canadian ethical regulations. Six- to ten-week-old female C57BL/6N mice (Charles River, 027) were used in experiments. One day prior to *S. boulardii* administration, groups of 2 to 5 mice were pretreated with a single dose of 20 mg of streptomycin as indicated. The following day, 5% sodium bicarbonate pretreatment, if indicated, was carried out 30 min prior to yeast gavage (*S. boulardii* bearing *ADH2*-m3F VHH or *ADH2* control HIV-2 VHH). Mice were administered 2×10^8^ CFU of the appropriate yeast suspension in either HEPES buffered saline (0.9% NaCl, 100 mM HEPES pH 7.5) or 5% sodium bicarbonate with HEPES buffer via oral gavage at the indicated frequency for each experiment. Fecal pellets were collected at the times indicated and homogenized in 1 mL PBS for 5 min at 30 rps (Retsch MM400) for determination of yeast CFU by serial dilution. At the experimental endpoint, ileum, cecum, and colon tissue, including luminal contents, were collected and homogenized in 1 mL PBS for 10 min at 30 rps. All homogenates were serially diluted and plated on YPD containing 200 μg/mL G418 and 100 μg/mL ampicillin for CFU determination.

### DSS colitis model

Two days following the initial gavage of *S. boulardii* strains or control buffer, mice were administered sterile 2% to 3% DSS (Thermo Fisher Scientific, CAS: 9011-18-1 Cat#J1448922) in double distilled water *ad libitum*. After 5 days, mice were shifted to regular drinking water and euthanized after having reached either 20% weight loss or experimental endpoint (up to 2 weeks post initial gavage). Fecal pellets and tissue were collected and processed as above. Disease activity index (DAI) was scored as described previously^70^. Briefly, DAI scores were determined by evaluating for body weight loss, stool consistency and occult/gross bleeding on the days indicated, as stratified in Supplementary Table 6. DIA scores are consistent with clinical symptoms observed in human inflammatory bowel disease. Statistical analyses were performed using GraphPad Prism v.98. Significance was tested by one-way ANOVA on ranks with Kruskal–Wallis test and Dunn’s post hoc analysis for multiple comparisons.

### 16S rDNA microbiome sequencing

Genomic DNA was extracted as described with modifications ^71^. Samples were transferred to screw cap tubes containing 2.8 mm ceramic beads, 0.1 mm glass beads, 5M guanidinium thiocyanate, 0.1M EDTA, 0.5% sarcosyl, and sodium phosphate buffer as described ^71^. Samples were homogenized with a bead beater, centrifuged, and the supernatant processed using a MagMAXExpress 96-Deep Well Magnetic Particle Processor (Applied Biosystems) with a Multi-Sample kit (Life Technologies, catalogue #4413022). Purified DNA was used to amplify variable regions 3 and 4 of the 16S rRNA gene by two-stage nested PCR. Initially samples were amplified in the 8f (AGAGTTTGATCCTGGCTCAG) to 926r (CCGTCAATTCCTTTRAGTTT) region of the 16S gene. The reaction was carried out at 94°C for 5 min, 15 cycles of 94°C for 30 sec, 56°C for 30 sec and 72°C for 60 sec, with a final extension of 72°C for 10 min. PCR products were used as template in the second stage of PCR in which the 341F (CCTACGGGNGGCWGCAG) to 806R (GGACTACNVGGGTWTCTAAT) region was amplified as previously described^72^. The reaction was carried out at 94°C for 5 min, 5 cycles at 94°C for 30 sec, 47°C for 30 sec and 72°C for 30 sec, followed by 25 cycles of 94°C for 30 seco, 50°C for 30 sec and 72°C for 30 sec, with a final extension of 72°C for 10 min. Resulting PCR products were visualized on a 1.5% agarose gel. Positive amplicons were normalized by pooling appropriately based on band intensity and sequenced on an Illumina MiSeq platform.

Bioinformatic analysis was performed as previously described^73–75^. Briefly, raw 16S amplicon sequences and metadata were demultiplexed using *split_libraries_fastq.*py script implemented in *QIIME2*^76^. Demultiplexed fastq files were split into sample-specific fastq files using *split_sequence_file_on_sample_ids*.py script from *QIIME2*. Individual fastq files without non-biological nucleotides were processed using Divisive Amplicon Denoising Algorithm (DADA) pipeline^77^. The output of the dada2 pipeline (feature table of amplicon sequence variants) was processed for alpha and beta diversity analysis using *phyloseq*^78^, and *microbiomeSeq* (http://www.github.com/umerijaz/microbiomeSeq) packages in R. We analyzed variance (ANOVA) among sample categories while measuring the of α-diversity measures using *plot_anova_diversity* function in *microbiomeSeq* package. Permutational multivariate analysis of variance (PERMANOVA) with 999 permutations was performed on all principal coordinates obtained during CCA with the *ordination* function of the *microbiomeSeq* package. The pairwise correlation was performed between the microbiome (genera) and metabolomics (metabolites) data using the *microbiomeSeq* package.

## Abbreviations

IBD: inflammatory bowel disease
TNF: tumor necrosis factor
VHH: variable heavy domain of heavy chain antibody
scFv: single-chain fragment variable
Fab: fragment antigen-binding region

## Author contributions

Roger Palou: methodology, formal analysis, investigation, writing - original draft, writing - review & editing, visualization; Almer van der Sloot: conceptualization, methodology, writing - original draft, writing - review & editing, supervision; Aline Fiebig: methodology, formal analysis, investigation, writing - review & editing, visualization; Megan Zangara: methodology, formal analysis, investigation; Naseer Sangwan: methodology, formal analysis; Maria Sanchez Osuna: methodology, formal analysis, investigation; Bushra Ilyas: methodology, investigation; Haley Zubyk: methodology, investigation; Michael Cook: investigation; Gerard D Wright: conceptualization, supervision, funding acquisition; Brian Coombes: conceptualization, writing - review & editing, supervision, funding acquisition; Mike Tyers: conceptualization, writing - original draft, writing - review & editing, supervision, funding acquisition

## Conflict of interest

The authors declare no conflict of interest.

## Acknowledgments

We thank Xingjian Xu for initial technical assistance, Jason Moffat and Dev Sidhu for antibody clones, and Bruce Futcher for plasmid pGZ110.

## Funding

This study was supported by grants from The W. Garfield Weston Foundation (to B.K.C., G.D.W. and M.T.), Genome Canada and Genome Quebec (to G.D.W. and M.T.), and the Canadian Institutes of Health Research (PJT-180315 to B.K.C.; FDN-167277 to M.T.).

## Supplementary Figure Legends

**Figure S1.**
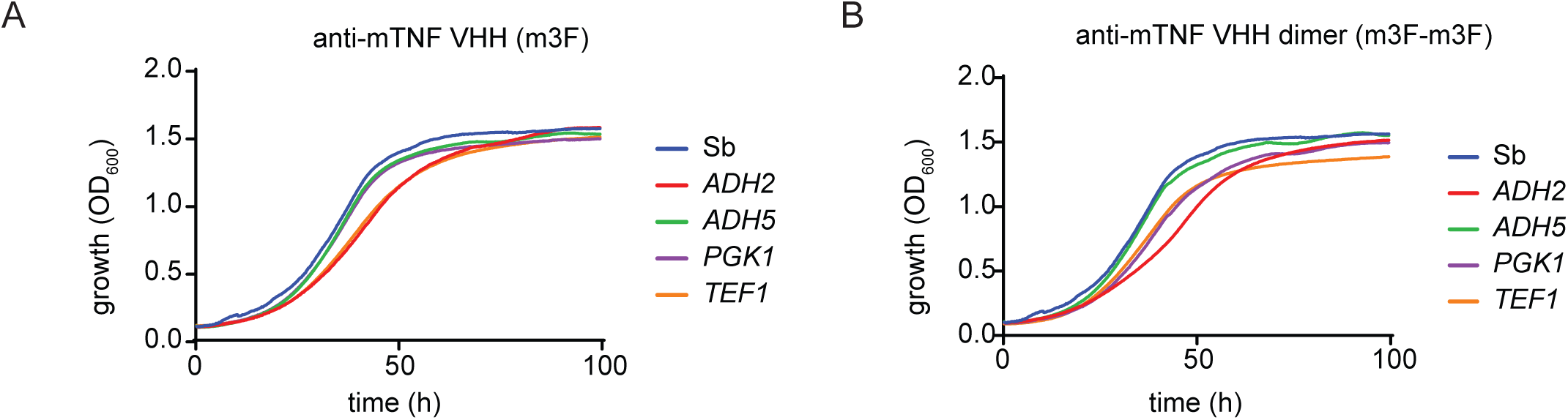
Effect of anti-mTNF VHH on *S. boulardii* proliferation *in vitro*. **A)** Growth rate of strains producing anti-mTNF VHH (m3F) produced under the control of *ADH2*, *ADH5*, *PGK1* and *TEF1* promoters were determined by OD_595_ in a shaker-reader at 37°C. **B)** Growth rate of strains producing anti-mTNF VHH tandem dimer (m3F-m3F) produced under the control of *ADH2*, *ADH5*, *PGK1* and *TEF1* promoters were determined by OD_595_ in a shaker-reader at 37°C. Wild type S. *boulardii* (Sb) was used as a control.

**Figure S2.**
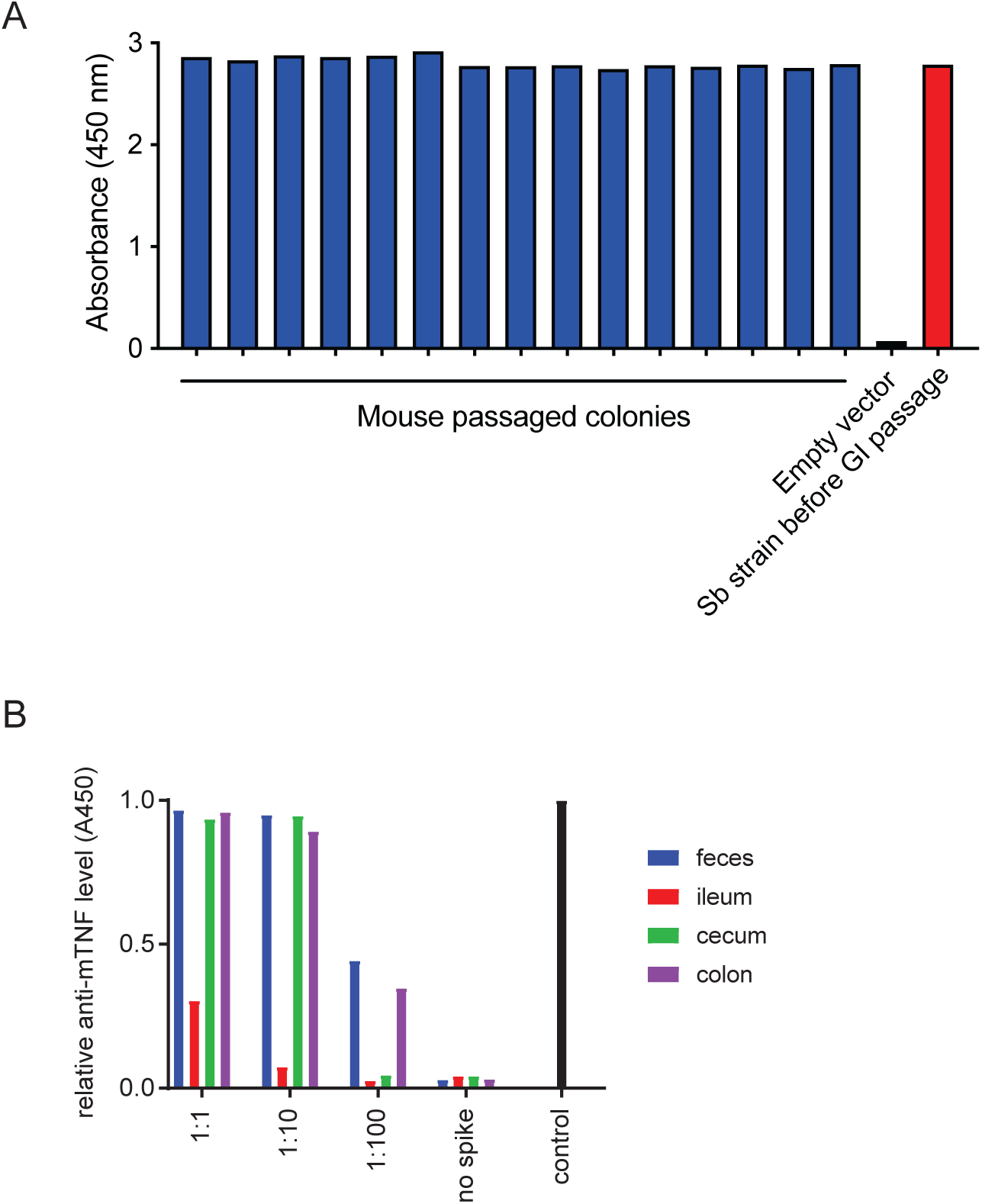
Production of anti-mTNF VHH by *S. boulardii* after passage through the mouse GI tract. **A)** Individual *S. boulardii* colonies obtained from feces were assessed for anti-mTNF VHH (m3F) expression by ELISA. **B)** Stability of anti**-**mTNF m3F VHH activity in indicated tissue and feces extracts. Dilutions of anti-mTNF VHH produced by *S. boulardii* in culture medium were incubated with tissue extracts and assessed for binding activity by ELISA.

**Figure S3.**
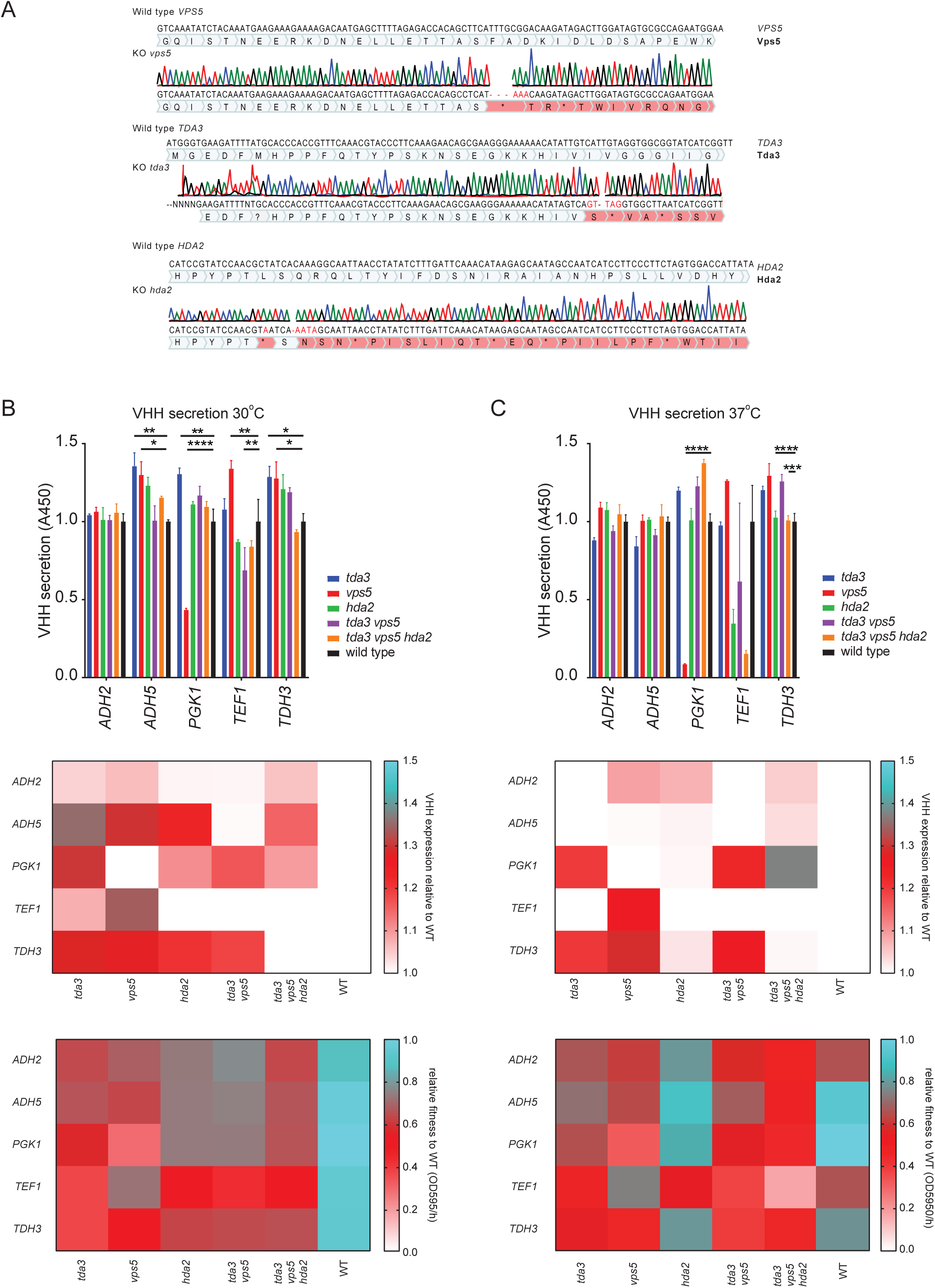
Effect of secretory pathway mutations on VHH production by *S. boulardii*. The secretory pathway genes *TDA3, VPA5* and *HDA2* were disrupted by CRISPR-mediated introduction of frameshift mutations. **A)** Sequence validation of mutant alleles. Sequences were rendered and visualized in Benchling. **B)** Single, double, and triple mutants were transformed with plasmids expressing anti-mTNF VHH (m3F) under the control of the *ADH2*, *ADH5*, *PGK1*, *TEF1* and *TDH3* promoters. Cells were grown in XY rich medium with 2% glycerol+2% ethanol at 30°C (*left*) in a shaker-reader for 3 days. VHH secretion was quantified by ELISA using immobilized anti-VHH antibody (top) and normalized to wild type *S. boulardii* expressing anti-mTNF m3F VHH from the *PGK1* promoter (middle). Mean ± SEM are indicated. Statistical analysis by Tukey’s ANOVA (* p<0.05, ** p<0.01, *** p<0.001, **** p <0.0001). Cell fitness was measured as the slope of the exponential growth phase for three independent colonies and normalized to wild type *S. boulardii* expressing anti-mTNF (m3F) VHH from the *ADH2* promoter (bottom). **C)** As for panel B but with cell culture at 37°C.

**Figure S4.**
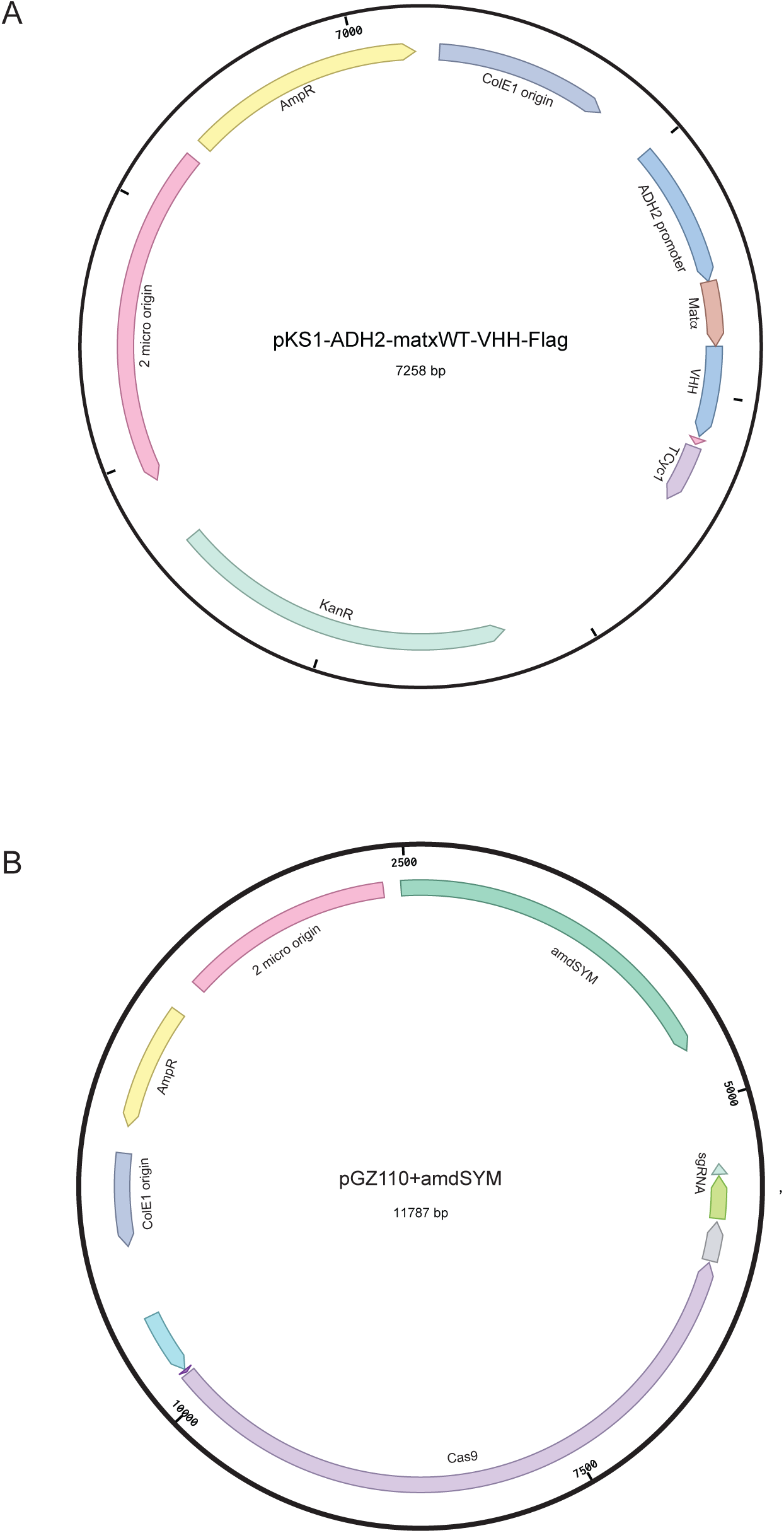
Plasmid map for constructs used in this study. **A)** pKS-p*ADH2*-*MAT*αWT-VHH-FLAG-*CYC1*term-KanMX for VHH expression and secretion. Variations of this construct replaced promoters, secretion signals and VHH coding regions. **B)** pGZ110-Cas9-amdSYM for *S. boulardii* genome engineering. Variations of this construct replaced sgRNA and repair template regions.

## Supporting information

**Supplementary Table 1.** List of secretion signals.

|  | Secretory Peptide Sequence | Origin |
| --- | --- | --- |
| Amylase | MVAWWSLFLYGLQVAAPALA | 79 |
| Glucanase | MQRPFLLAYLVLSLLFNSAL | 80 |
| Inulase | MKFAYSLLLPLAGVSA | 81 |
| Invertase | MLLQAFLFLLAGFAAKISA | 82 |
| Lysozyme | MRSLLILVLCFLPLAALG | 82 |
| Matα WT | MRFPSIFTAVLFAASSALAAPVNTTTEDETAQIPAEAVIGYLDLEGDFDVAVLFPNSTNNG<br>LLFINTTASIAAKEEGVQLDKR | |
| Matα App8<br>(Ala22) | MRFPSIFTAVLFAASSALAAPANTTTEDETAQIPAEAVIDYSDLEGDFDAALPLSNSTNNG<br>LSSTNTTASIAAKEEGVQLDKR | 32 |
| Matα App8 | MRFPSIFTAVLFAASSALAAPVNTTTEDETAQIPAEAVIDYSDLEGDFDAALPLSNSTNNG | 32 |
| A22V | LSSTNTTASIAAKEEGVQLDKR |  |
| BBa_K416<br>003<br>(BBaK) | MKVLIVLLAIFAALPLALAQPVISTTVGSAAEGSLDKR | 33 |

\* mutated residues in red with respect to the wild type Matα signal

**Supplementary Table 2.** List of VHH nanobody sequences.

| Name | Code | Target | Sequence | Origin |
| --- | --- | --- | --- | --- |
| HIV-1 | L8Cj3 | HIV-1 | VQLVESGGGLVQAGGFLRLSCELRGSI FNQYAMAWFRQAPG<br>KREFVAGMGAVPHYGEFVKGRFTISRDN AKSTVYLQMSSL<br>KPEDTAIFYCARSKSTYISYNSNGYDYWGRGTQVTVSSGGSD<br>YKDDDDK | 40 |
| HIV-2 | J3r | HIV-1 | VQLQESGGGLVQAGGSLRLSCELRGSI FNQYAMAWFRQAPG<br>KREFVAGMGAVPHYGEFVKGRFTISRDN AKSTVYLQMNSL<br>KPEDTAIFYCARSKSTYISYNSNGYDYWGQGTQVTVSSGGSD<br>YKDDDDK | 40 |
| HIV-3 | L911F1<br>F | HIV-1 | VQLVESGGALVQAGRSRLRLSCAASGNAFTIDAAAWYRQAPG<br>KQREPVATILSGGTTNYADSVKGRFTISRDNVKNTVYLQMNS<br>LKPEDTAVYYCYVPMVYYSGRYNDVWGQGTQVTVSSGGSD<br>YKDDDDK | 40 |
| CD-1 | A4.2 | TcdA | EGVQLDKREAQVKLEESGGGLVQAGGSLRLSCAASGRFTFNT<br>LSMGWFRQAPGKREFVAAVSRSGGSTYYADSVKGRFTISRDN<br>NAKNTVYLQMNSLKPEDTAVYYCAAATKSNTTAYRLSFDY<br>WGQGTQVTVSSGGSDYKDDDD | 83 |
| CD-2 | A20.1 | TcdA | EGVQLDKREAQVQLVESGGGLAQAGGSLRLSCAASGRFTFSM<br>DPMWFRQPPGKREFVAAAGSSTGRTTYADSVKGRFTISRDN<br>NAKNTVYLQMNSLKPEDTAVYYCAAAPYGANWYRDEYDY<br>WGQGTQVTVSSGGSDYKDDDD | 83 |
| CD-3 | A26.8 | TcdA | EGVQLDKREAQVKLEESGGGLVQAGGSLRLSCAASERTFSRY<br>PVAWFRQAPGAEREFVAVISSTGTSTYYADSVKGRFTISRDN<br>AKVTVYLQMNNLKREDTAVYFCVNSQRTRLQDPNEYDYW<br>GQGTQVTVSSGGSDYKDDDD | 83 |
| CD-4 | A5.1 | TcdA | EGVQLDKREAQVKLEESGGGLVQAGGSLRLSCAASGRFTFSM<br>YRMGWFRQAPGKREFVGVITRNGSSTYYADSVKGRFTISRDN<br>NAKNTVYLQMNSLKPEDTALYYCAATSGSSYLDAAHVYDY<br>WGQGTQVTVSSGGSDYKDDDD | 83 |
| CD-5 | 4NC2 | TcdB | EGVQLDKREAQVQLVESGGGLVQAGGSLRLSCAASGLTFSR<br>YVMGWFRQAPGKREFVAAITWGGTPNYADSVKGRFTISRDN<br>NSKNTQYLMNSLKPEDTAVYYCAAGLGWDSRYSQSYNYW | 84 |
|  |  |  | GQGTQVTVSSGGSDYKDDDD |  |
| CJ-1 | V1Flag | *C. jejuni* | EGVQLDKREAQVKLEESGGGLVQAGGSRRLSCATSGLTFRNF<br>HMAWFRQVAGKEREVVA AISWSRDRQYYPDPVKGRFTITRD<br>NAKNTVYLQMNSLKPEDTAVYYCAARTASASGDWYKGSYQ<br>YWGQGTQVTVSSGGSDYKDDDD | 85 |
| CJ-2 | V6Flag | *C. jejuni* | EGVQLDKREAQVKLEESGGGLVQAGGSLRVSTASVSTFSIN<br>ALGWYRQAPGKARELVAAIGSDGTVYYTDSVKGRFTISRDN<br>AKNTVSLQMSSLKPEDTAVYYCNAAGKRIGSDGSIWFAVASF<br>GSWGQGTQVTVSSGGSDYKDDDD | 85 |
| ETEC-1 | K922 | F4 fimbriae | EGVQLDKREAQVQLQESGGGLVQPGGSLRLSCLVSGGTFSW<br>YAMGWFRQAPGKEREFVATVSRGGGSSYYADSVKGRFTISR<br>DNAKNTVYLQMNSLKPEDTAVYYCAAGRGAPSDTGRPDEY<br>DYWGQGTQVTVSSGGSDYKDDDD | 86 |
| ETEC-2 | 4PC2 | Stx2 | EGVQLDKREAQVQLQESGGGLVQAGGSLRLSCAVSGSIFRLS<br>TMGWYRQAPGKQREFVASITSYGDTNYRDSVKGRFTISRDN<br>AKNTVYLQMNSLKPEDTAVYYCNANIEAGTYYPGRDYGW<br>QGTQVTVSSGGSDYKDDDD | 87 |
| LM-1 | VH303 | *L. monocytogenes* | EGVQLDKREAQVQLVESGGGLVQPGRSLRLSCAASGHTYSTY<br>CMGWVRRAPGKGEELVARINVGGSSWTYADSVKGRFTISAD<br>TSKNTAYLQMNSLRAEDTAVYYCTLHRFCNTWSLGTNLNYS<br>GQGLTVTVSSGGSDYKDDDD |  |
| LM-2 | VH-1bag | *L. monocytogenes* | EVQLVESGGGLVQPGRSLRLSCAASGFNIKDTYIGWVRRAPG<br>KGEELVARIYPTNGYTRYADSVKGRFTISADTSKNTAYLQMN<br>SLRAEDTAVYYCARWGGDGFYAMDYWGQGLTVTVSSGGSD<br>YKDDDD |  |
| hTNF-1 | 5M2I | human TNF | EGVQLDKREAQVQLVESGGGLVQAGGSLSLSCASGRSLSNY<br>YMGWFRQAPGKERELLGNISWRGYNIYYKDSVKGRFTISR<br>DAKNTIYLMNRLKPEDTAVYYCAASILPLSDDPGWNTYWG<br>QGTQVTVSSGGSDYKDDDD | 45 |
| hTNF-2 | 5M2J | human TNF | EGVQLDKREAQVQLVESGGGLVQPGGSLRLSCAASGFTFSNY<br>WMYWVWRQAPGKLEWVSEINTNGLITKYPDSVKGRFTISR<br>NAKNTLYLMNLSLKPEDTALYYCARSPSGFNRGQGTQVTVS<br>SGGSDYKDDDD | 45 |
| hTNF-3 | 5M2M | human TNF | EGVQLDKREAQVQLQESGGGLVQPGGSLRLSCAASGRFTSD<br>HSGYTYTIGWFRQAPGKEREFVARIYWSSGNTYYADSVKGRF<br>AISRDIAKNTVDLTMNNLEPEDTAVYYCAARDGIPTSRSVESY<br>NYWGQGTQVTVSSGGSDYKDDDD | 45 |
| hTNF-4 | 5M2M-5M2I | human TNF | EGVQLDKREAQVQLQESGGGLVQPGGSLRLSCAASGRFTSD<br>HSGYTYTIGWFRQAPGKEREFVARIYWSSGNTYYADSVKGRF<br>AISRDIAKNTVDLTMNNLEPEDTAVYYCAARDGIPTSRSVESY<br>NYWGQGTQVTVGGSGGGSGGGSQVQLVESGGGLVQAGGS | 45 |
|  |  |  | LSLSCSASGRSLSNYYMGWFRQAPGKERELLGNISWRGYNIY<br>YKDSVKGRFTISRDDAKNTIYLQMNRLKPEDTAVYYCAASIL<br>PLSDDPGWNTYWGQGTQVTVSSGGSDYKDDDD |  |
| mTNF | M3F | mouse TNF | QVQLQDSGGGLVQAGGSLRLSCAASGGTFSSIIMAWFRQAPG<br>KEREFVGAVSWSGGTTVYADSVLGRFEISRDSARKSVYLQM<br>NSLKPEDTAVYYCAARPYQKYNWASASYNVWGQGTQVTVS<br>SEPKTPKPQPTVSSGGSDYKDDDDK | 46 |

**Supplementary Table 3.**
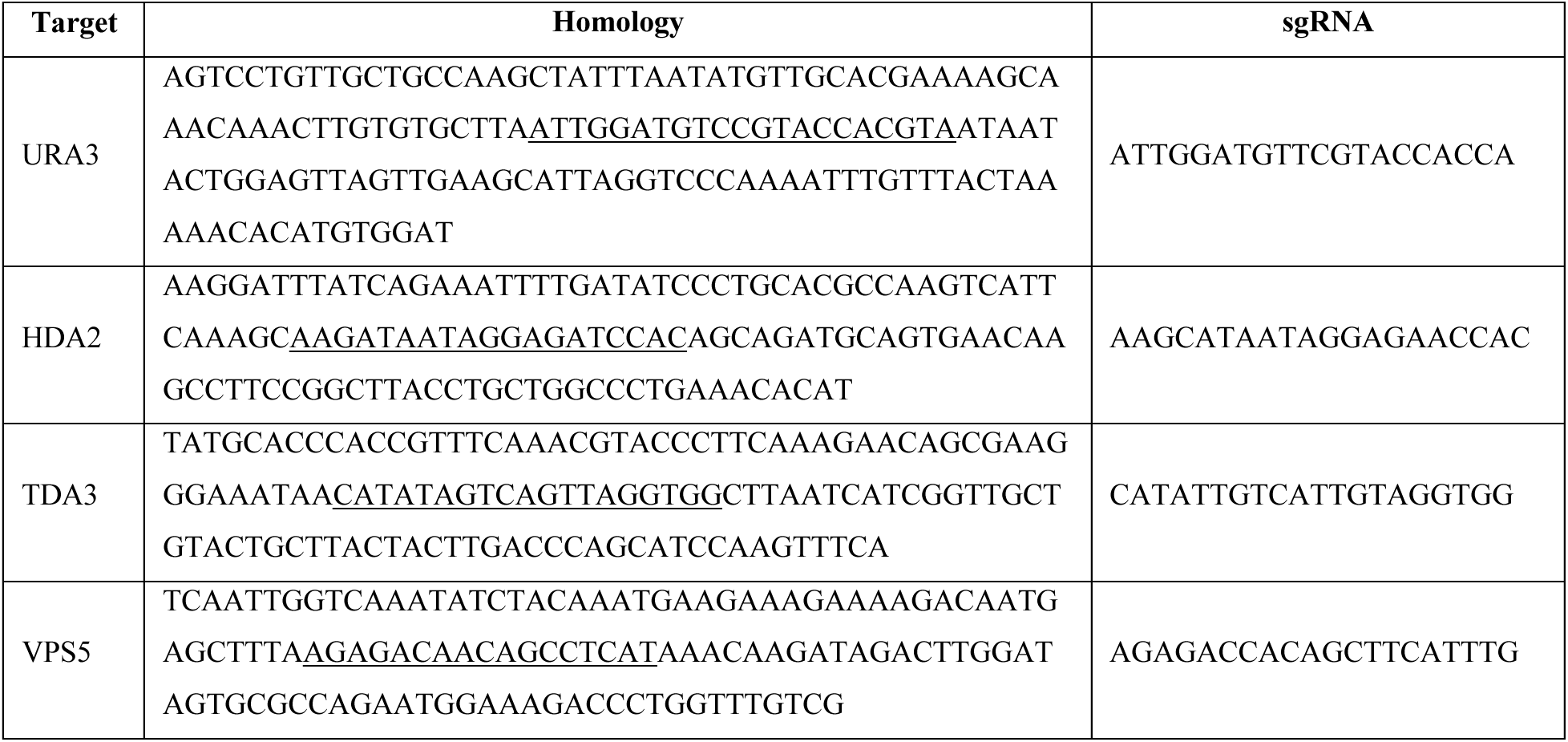
List of homology repair and sgRNA sequences.

**Supplementary Table 4.** Plasmids used in this study.

| ID | Plasmid Structure | Source |
| --- | --- | --- |
| MT4954 | pKS1ST | DualSystems Biotech |
| MT4651 | pD1204-GAL1-MATαWT-[CDTb-B12-Light chain, Fab']-FLAG-TCyc1-URA3 | This study |
| MT4652 | pD1204-GAL1-MATαAla22-[CDTb-B12-Light chain, Fab']-FLAG-TCyc1-URA3 | This study |
| MT4653 | pD1204-GAL1-MATαVal22-[CDTb-B12-Light chain, Fab']-FLAG-TCyc1-URA3 | This study |
| MT4654 | pD1204-GAL1-BBaK-[CDTb-B12-Light chain, Fab']-FLAG-TCyc1-URA3 | This study |
| MT4655 | pD1204-GAL1-Lysozyme-[CDTb-B12-Light chain, Fab']-FLAG-TCyc1-URA3 | This study |
| MT4656 | pD1204-GAL1-Invertase-[CDTb-B12-Light chain, Fab']-FLAG-TCyc1-URA3 | This study |
| MT4657 | pD1204-GAL1-Glucanase-[CDTb-B12-Light chain, Fab']-FLAG-TCyc1-URA3 | This study |
| MT4658 | pD1204-GAL1-Amylase-[CDTb-B12-Light chain, Fab']-FLAG-TCyc1-URA3 | This study |
| MT4659 | pD1204-GAL1-Inulase-[CDTb-B12-Light chain, Fab']-FLAG-TCyc1-URA3 | This study |
| MT4660 | pD1204-GAL1-MATαWT-scFvGC132-FLAG-TCyc1-URA3 | This study |
| MT4661 | pD1204-GAL1-MATαWT-LaG16 VHH-FLAG-TCyc1-URA3 | This study |
| MT4663 | pGZ110-Cas9-amdSYM | This study |
| MT4664 | pGZ110-Cas9-amdSYM-URA3sgRNA | This study |
| MT4665 | pKS-pADH2- MATαWT -J3r-FLAG-TCyc1-KanMX | This study |
| MT4666 | pKS-pADH2- MATαWT -L8Cj3-FLAG-TCyc1-KanMX | This study |
| MT4667 | pKS-pADH2- MATαWT -L911F1F-FLAG-TCyc1-KanMX | This study |
| MT4668 | pKS-pADH2- MATαWT -A4.2-FLAG-TCyc1-KanMX | This study |
| MT4669 | pKS-pADH2- MATαWT -A20.1-FLAG-TCyc1-KanMX | This study |
| MT4670 | pKS-pADH2- MATαWT -A26.8-FLAG-TCyc1-KanMX | This study |
| MT4671 | pKS-pADH2- MATαWT -4NC2-FLAG-TCyc1-KanMX | This study |
| MT4672 | pKS-pADH2- MATαWT -FlagV1-FLAG-TCyc1-KanMX | This study |
| MT4673 | pKS-pADH2- MATαWT -FlagV6-FLAG-TCyc1-KanMX | This study |
| MT4674 | pKS-pADH2- MATαWT -K922-FLAG-TCyc1-KanMX | This study |
| MT4675 | pKS-pADH2- MATαWT -4PC2-FLAG-TCyc1-KanMX | This study |
| MT4676 | pKS-pADH2- MATαWT -5M2I-FLAG-TCyc1-KanMX | This study |
| MT4677 | pKS-pADH2- MATαWT -5M2J-FLAG-TCyc1-KanMX | This study |
| MT4678 | pKS-pADH2- MATαWT -5M2M-FLAG-TCyc1-KanMX | This study |
| MT4681 | pKS-pADH2- MATαWT -VH303-FLAG-TCyc1-KanMX | This study |
| MT4682 | pKS-pADH2- MATαWT -VH-1bag-FLAG-TCyc1-KanMX | This study |
| MT4683 | pKS-pADH2- MATαWT -5M2M-5M2I-FLAG-TCyc1-KanMX | This study |
| MT4684 | pKS-pADH2- MATαWT -m3F-FLAG-TCyc1-KanMX | This study |
| MT4685 | pKS-pADH2- MATαWT -m3F-6xHIS-TCyc1-KanMX | This study |
| MT4686 | pKS-pADH2- MATαWT -m3F-m3F-FLAG-TCyc1-KanMX | This study |
| MT4687 | pKS-pADH5- MATαWT -m3F-FLAG-TCyc1-KanMX | This study |
| MT4688 | pKS-pPGK1- MATαWT -m3F-FLAG-TCyc1-KanMX | This study |
| MT4689 | pKS-pTDH3- MATαWT -m3F-FLAG-TCyc1-KanMX | This study |
| MT4690 | pKS-pTEF1- MATαWT -m3F-FLAG-TCyc1-KanMX | This study |
| MT4691 | pKS-pADH5- MATαWT -m3F-m3F-FLAG-TCyc1-KanMX | This study |
| MT4692 | pKS-pPGK1- MATαWT -m3F-m3F-FLAG-TCyc1-KanMX | This study |
| MT4693 | pKS-pTEF1- MATαWT -m3F-m3F-FLAG-TCyc1-KanMX | This study |
| MT4697 | pKS-pGAL1- MATαWT -5M2I-6xHIS-TCyc1-KanMX | This study |
| MT4698 | pKS-pGAL1- MATαWT -5M2I-6xHIS-TCyc1-KanMX | This study |
| MT4699 | pKS-pGAL1- MATαWT -5M2M-5M2I-6xHIS-TCyc1-KanMX | This study |
| MT4700 | pKS-ppADH2- MATαWT -J3r-6xHIS-TCyc1-KanMX | This study |
| MT4701 | pKS-pADH2- MATαWT -A5.1-FLAG-KanMX | This study |

**Supplementary Table 5.** Strains used in this study.

| ID | Yeast | Genotype | Origin |
| --- | --- | --- | --- |
| MTy5073 | *S. boulardii* | MYA-796 | ATCC |
| MTy1580 | *S. cerevisiae* | Sigma1278b | Tyers lab |
| MTy5084 | *S. boulardii* | MYA-796 *ura3* | This study |
| MTy5087 | *S. boulardii* | MYA-796 *hda2* | This study |
| MTy5088 | *S. boulardii* | MYA-796 *tda3* | This study |
| MTy5089 | *S. boulardii* | MYA-796 *vps5* | This study |
| MTy5090 | *S. boulardii* | MYA-796 *tda3 vps5* | This study |
| MTy5091 | *S. boulardii* | MYA-796 *hda2 vps5 tda3* | This study |

**Supplementary Table 6:**
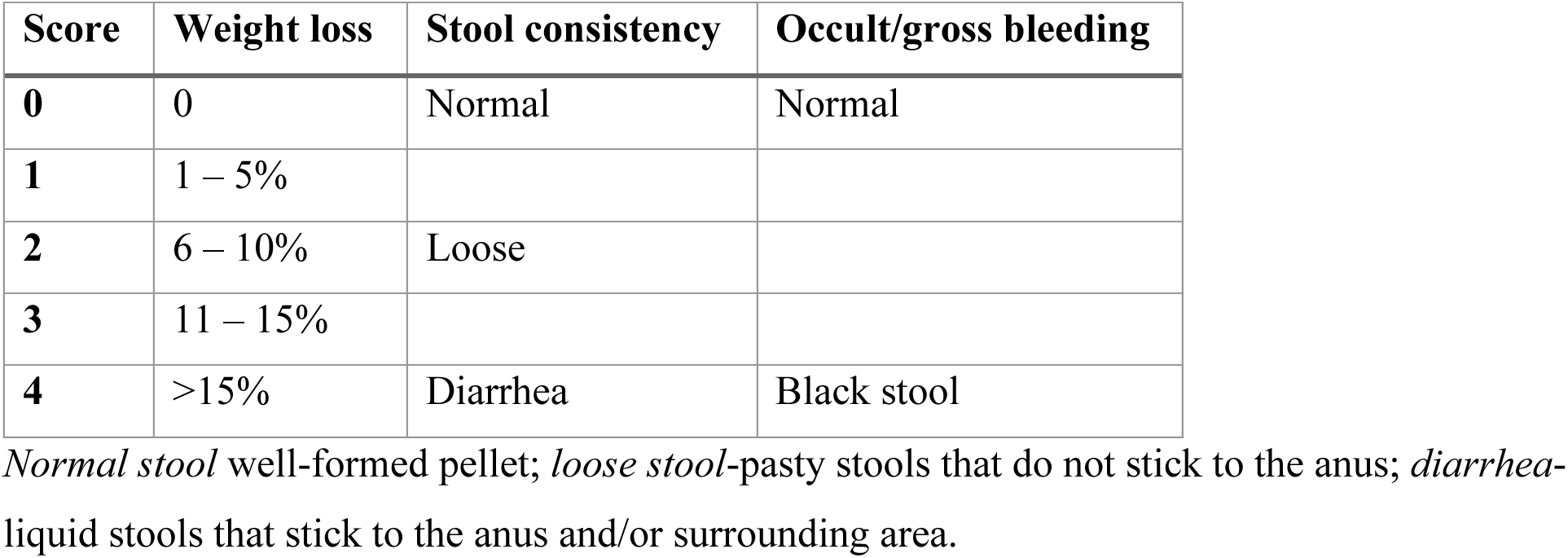
Disease activity index score scale.

## Notes

### Competing Interest Statement

The authors have declared no competing interest.

